# Robustness of the inference of the distribution of fitness effects to selective sweeps

**DOI:** 10.64898/2026.10.25.761705

**Authors:** Emma E. Wade, Kirk E. Lohmueller

**Affiliations:** Bioinformatics Interdepartmental Program, University of California, Los Angeles; Department of Ecology and Evolutionary Biology, University of California, Los Angeles; Department of Human Genetics, University of California, Los Angeles

## Abstract

The distribution of fitness effects (DFE) describes the proportion of new mutations with certain selective strengths. It is a crucial quantity for the study of complex disease and evolutionary genetics. Population genetics approaches infer the DFE using the site frequency spectrum (SFS), which summarizes allele frequencies within a sample. These approaches first fit a demographic model to an SFS of putatively neutral sites, then infer the DFE from an SFS of selected sites conditional on that demographic model. This approach assumes that sites are independent and that population size changes affect selected and neutral sites equally. Selective sweeps may violate these assumptions. As a beneficial mutation rises in frequency, nearby sites hitchhike alongside the beneficial mutation. This distortion might differ between neutral and selected sites.. Here, we evaluate the performance of the DFE inference method, fit∂*a*∂*i*, in the presence of selective sweeps. We simulate genomic segments containing neutral and deleterious mutations drawn from a known DFE under varying selective sweep conditions, and test whether fit∂*a*∂*i* can recover the true DFE with a flexible demographic model. When SFS are sampled immediately after fixation of a beneficial mutation, the demographic models absorb the sweep’s effect as a population expansion, and the DFE is recovered relatively well. Performance declines at lower recombination rates and when using the unfolded SFS, leading to an overestimation of the strength of purifying selection. We then model recurrent selective sweeps at rates consistent with human polymorphism and divergence data. The demographic model captures these distortions, and the DFE is unbiased. Our results suggest that DFE inferences are unlikely to be substantially biased by selective sweeps, but that future studies should consider using the folded SFS and excluding regions of low recombination rate.

## Introduction

Mutations vary in their effects on fitness, ranging from deleterious to neutral to beneficial. The distribution of fitness effects (DFE) describes this continuum of selection coefficients of new mutations within a population [1]. Accurate estimation of the DFE is critical as it shapes the genetic architecture of complex traits [2], the molecular clock [3], evolution of recombination [4], and genetic rescue [5–7].

Population genetic approaches for DFE inference, including fit∂*a*∂*i[8]*, leverage that a mutation’s fitness effect influences its frequency [9–12]. These methods summarize allele frequency across many sites in the site frequency spectrum (SFS) and use it to estimate the proportion of new mutations with certain selective strengths. Classically, the Poisson random field (PRF) model on which SFS-based DFE inference methods are based assumes that all mutations are independent of one another. As such, the model does not directly model linked selection, which can impact surrounding patterns of variation [13–16]. In addition to the direct effects of natural selection, SFS are also affected by population size history [17]. To account for these confounders, SFS-based DFE inferences are divided into two steps. First, a demographic model is fit to a SFS composed of putatively neutral sites, often synonymous sites, which are assumed to only be affected by demography and linked selection. Second, a DFE is fit to selected sites, usually nonsynonymous mutations, conditional on this demographic model. This approach assumes that population size changes and linked selection affect synonymous and nonsynonymous variants similarly and that the demographic model is able to absorb these effects, leaving the DFE unbiased. Intuitively, while the PRF model does not model linked selection directly, if linked selection affects genealogies of nearby neutral sites, then the demographic history inferred at these neutral sites captures both the effects of demography and linked selection. As such, the correction provides some degree of correction for the unmodeled linked selection.

Indeed, using the demographic correction described above, fit∂*a*∂*i* appears robust to the reduction in genetic diversity caused by linked purifying selection (background selection) [8,18,19]. However, it is less clear whether DFE inferences are biased by linked positive selection. When a beneficial mutation rises to high frequency, a process called a selective sweep, nearby alleles “hitchhike” alongside the beneficial mutation [20]. This skews the SFS towards rare and high frequency derived alleles [21,22].

Evidence suggests positive selection affects selected sites differently than neutral sites, meaning a demographic model fit to synonymous sites may be insufficient to control for genetic hitchhiking distortions in the nonsynonymous SFS. When selected sites are linked, they interfere with each other and reduce the efficacy of natural selection, a phenomenon called Hill-Robertson Interference (HRI) [23,24] which is strongest in regions of low recombination [25]. During a sweep, a beneficial mutation can carry linked weakly deleterious variation to high frequencies [26,27], consistent with higher linkage disequilibrium (LD) between common nonsynonymous variants than synonymous variants during a sweep [28,29]. Conversely, strongly deleterious variants may impede genetic hitchhiking [30]. Because these interactions do not occur at synonymous sites, a demographic model fit to synonymous sites cannot absorb them.

Messer and Petrov (2013) [31] examined whether segregating beneficial mutations caused biases in DFE-alpha [10], which first infers a simple demographic history (constant or one size change) from synonymous sites and then fits a DFE to nonsynonymous sites conditional on that demography. In forward simulations of a human-like genome architecture with recurrent beneficial mutations and background selection, positive selection distorted both the selected and neutral SFSs which the demographic inference interpreted as a population expansion. Conditioned on this demographic history, the mean strength of purifying selection was overestimated. Tataru et al. (2017) also demonstrated this overestimation when not accounting for beneficial mutations in DFE-alpha as well as in polyDFE, which uses nuisance parameters instead of a demographic model to adjust for biases shared by both neutral and selected sites [32]. It has not been evaluated whether fit∂a∂i which can use more flexible demographic models may improve these biases. The extent to which these biases may be prevalent in real inferences is also unclear.

In this study, we run extensive forward-in-time simulations to test whether selective sweeps bias DFE inferences from fit∂a∂i. We first simulate genetic segments containing neutral and deleterious variation drawn from a known DFE, with a human-like and reduced recombination rates. We place a single beneficial mutation within these segments then terminate the simulations upon fixation, approximating the maximum amount of SFS distortion caused by a sweep. Using these simulations, we test whether fit∂a∂i, conditional on a flexible demographic model, can accurately infer the simulated deleterious DFE. Next, we test DFE inferences in simulations with recurrent beneficial mutations occurring at a rate expected by divergence and polymorphism data [33]. We find that single sweep caused a slight overestimation of the proportion strongly deleterious mutations particularly in regions of low recombination (r = 1 × 10^−9^ per bp per generation). Under a recurrent model, inferences did not differ from the true simulation proportions. These results suggest that DFE inference is largely reliable even in populations with frequent adaptation, but that estimates from low-recombination regions should be interpreted with caution.

## Methods

### Simulation overview

We first designed simulations containing a single sweep conditional-on-fixation, approximating the maximum SFS distortion from a selective sweep. We then run simulations with recurrent beneficial mutations to see if biases in the single sweep conditions generalize to a more biologically realistic scenario.

#### Single selective sweep conditional-on-fixation

Simulations were performed with the forward-in-time simulation framework, *SLiM5 [34]*. Each simulated segment contained 10 Mb of sequence. The population size was kept at a constant *N*=10,000 diploid individuals.

Each segment was randomly composed. Intergenic gaps were drawn from a uniform distribution between (500, 50,000) bp (mean ∼25 kb). Recombination events could occur within these gaps but they are not genomic elements, so mutations did not occur within them. Each gene started with an exon, then added intron-exon pairs. After each pair, another was added with probability 0.8. Exon and intron lengths were drawn from independent log normal distributions with mean ln(50) and ln(100), respectively. This resulted in an average composition of 1.47% exonic and 2.25% intronic sequence per segment.

Mutations entered the simulation at a rate of µ = 1. 5 × 10^−8^ per bp per generation. 69.8% of exonic mutations were nonsynonymous. The remaining 30.2% were synonymous. The selection coefficients, *s*, of nonsynonymous mutations were drawn from the gamma distributed human DFE estimated by Kim et al. 2017 (mean= -0.0131, shape=0.186)[8]. Intronic and synonymous exonic mutations were neutrally evolving (*s* = 0). All mutations were additive (*h* = 0. 5).

After a burn-in of 100,000 generations (10 × *N*), we placed a single beneficial mutation in the center of the simulated segment in one randomly chosen haplosome. Positive selection coefficients, *s*_*ben*_, of 0 (neutral control), 0.01, and 0.05 were tested. Neutral control simulations, *s* = 0, were continued for 2400 generations after the burn in, roughly the expected fixation time for the beneficial mutations with *s*_*ben*_ = 0. 01[35]. Simulations with *s* _*ben*_ > 0 were run conditional-on-fixation. In other words, the simulation continued until the beneficial mutation was fixed and then immediately terminated. If a beneficial mutation was lost, the simulation restarted post-burn in, at generation 100,000, with a new seed.

After the simulation was completed, a sample of *n* ∈ {100, 1000} haplosomes were drawn. SFS were recorded within windows extending 1, 3, and 5 Mb on either side of the positively selected site. Thus the SFS were pulled from sequence length totaling 2Mb, 6Mb, 10Mb, respectively. Exonic synonymous mutations and intronic mutations were combined to make the synonymous SFS. SFS of 22 independent SLiM simulated segments were summed together to form one replicate, so each replicate consisted of 22 completed sweeps. A total of 20 replicates were simulated for each unique parameter regime.

#### Recurrent selective sweeps

To simulate recurrent selective sweeps, we modified the selection coefficient of a proportion of nonsynonymous exonic mutations, *p*_*ben*_, to be beneficial with selective strength *s*_*ben*_ . The selection coefficients of the remaining nonsynonymous mutations, 1 − *p*_*ben*_, were drawn from the Kim et al. (2017) human DFE[8]. We tested values of *p*_*ben*_ and *s*_*ben*_ consistent with observed levels of polymorphism and divergence in humans (*log*_10_ (*s*_*ben*_ × *p*_*ben*_) =− 6) (Table 1) [33].

**Table 1.**
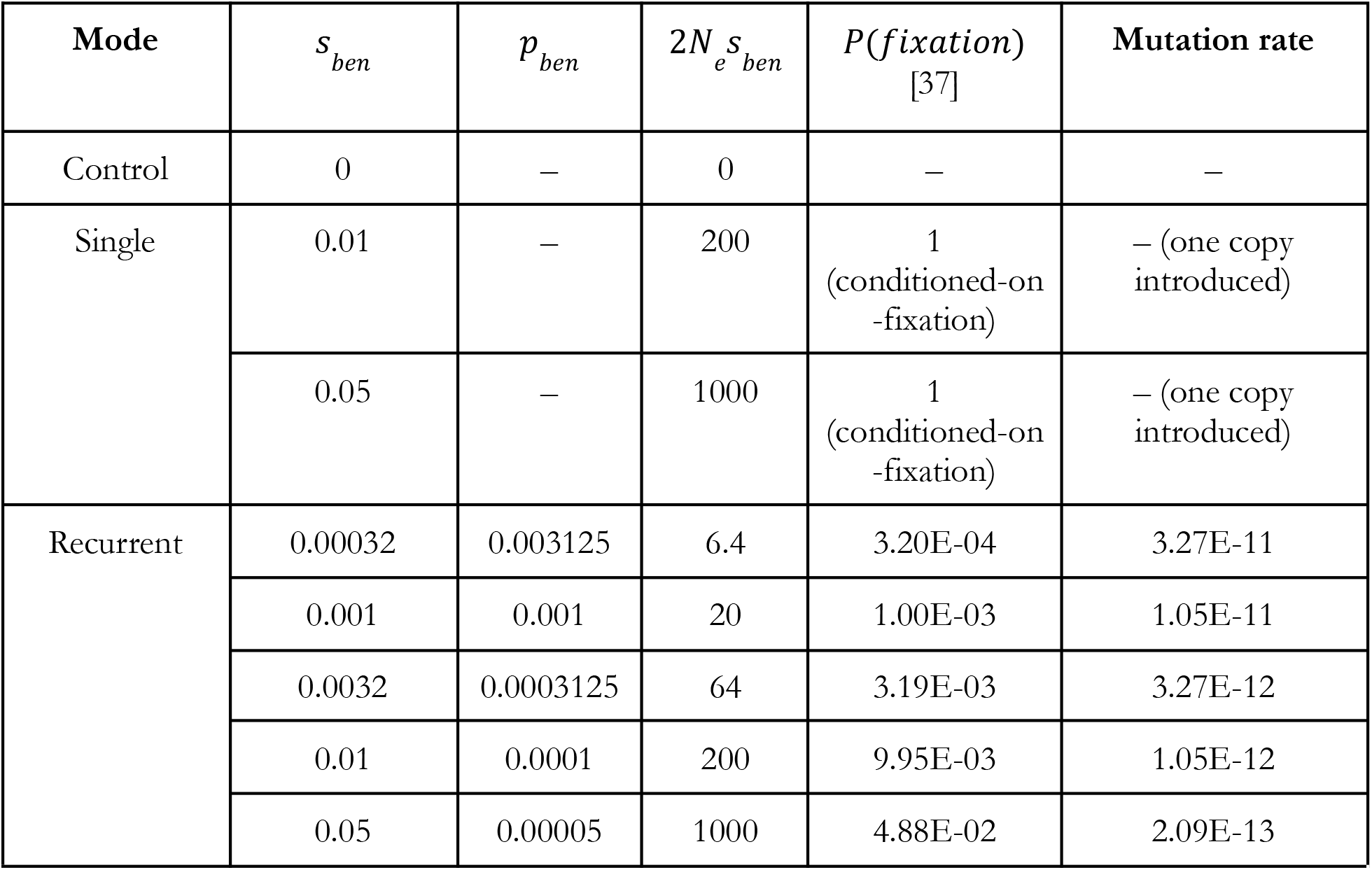
Sweep simulation conditions. The table lists the single-sweep and recurrent-sweep simulation regimes, with the probability of fixation and mutation rate for the beneficial mutations. *s*_*ben*_ is the unscaled selection coefficient of beneficial mutations in SLiM units (1+*hs*). *p*_*ben*_ is the proportion of new nonsynonymous mutations that are beneficial. The rest (1-*p*_*ben*_) are drawn from the Kim et al. (2017) DFE. *P*(*fixation*) is the fixation probability of the beneficial mutations. For single sweeps, it is 1 because each run is conditioned-on-fixation. For recurrent sweeps, Kimura’s probability for a new additive mutation is applied. The beneficial mutation rate is µ × 0. 698 × *p*_*ben*_ per exonic site per generation.

| Mode | $s_{ben}$ | $p_{ben}$ | $2N_e s_{ben}$ | $P(fixation)$<br>[37] | Mutation rate |
| --- | --- | --- | --- | --- | --- |
| Control | 0 | – | 0 | – | – |
| Single | 0.01 | – | 200 | 1<br>(conditioned-on<br>-fixation) | – (one copy<br>introduced) |
|  | 0.05 | – | 1000 | 1<br>(conditioned-on<br>-fixation) | – (one copy<br>introduced) |
| Recurrent | 0.00032 | 0.003125 | 6.4 | 3.20E-04 | 3.27E-11 |
|  | 0.001 | 0.001 | 20 | 1.00E-03 | 1.05E-11 |
|  | 0.0032 | 0.0003125 | 64 | 3.19E-03 | 3.27E-12 |
|  | 0.01 | 0.0001 | 200 | 9.95E-03 | 1.05E-12 |
|  | 0.05 | 0.00005 | 1000 | 4.88E-02 | 2.09E-13 |

Positive selection occurred for the full simulation duration, 102,400 generations, matching the number of generations in the neutral single sweep controls. The nonsynonymous SFS included all segregating nonsynonymous exonic mutations, both beneficial and deleterious. SFS were drawn from sample sizes of *n* ∈ {100, 1000} and recorded only for the full 10 Mb segment.

#### Simulations with varying recombination rate

Low recombination increases linkage disequilibrium (LD), enhancing HRI. Thus we inferred the DFE from simulations with the average recombination rates in the human genome *r* = 1 × 10^−8^ crossovers per site per generation [36] as well as a 10x decreased recombination rate, *r* = 1 × 10^−9^ crossovers per site per generation.

### Demographic inference

We used the diffusion-based approach ∂*a*∂*i* to identify the best fitting demographic model to our synonymous SFS [38,39]. As we are interested in whether a complex demographic model is able to capture the effects of genetic hitchhiking, and not in testing the accuracy of demographic inference to capture the true population size history (which is always set constant here), we fit demographic models with size changes. We first utilized a 2-epoch model where a single size change occurred at time, *T*, that changed the population size, ν, relative to *N*_*anc*_. To test if additional parametersimproved model fit, we fit 3-epoch models where two size changes occurred with parameters ν_*B*_, ν_*F*_,*T*_*B*_, and *T*_*F*_ .

In short, parameter optimization is done by initializing 33 randomly perturbed starting parameter values. We omitted population contraction starting values (ν < 1) as these are unlikely to lead to the best fitting model and increased runtime. Parameter values are repeatedly updated with L-BFGS-B for a maximum of 25 iterations. A fixed grid of 1200, 1400, 1600 pts was used for Richard extrapolation. We set parameter bounds, [0. 01 ≤ ν < ∞) and [1 × 10^−5^ ≤ *T* ≤ 10], to speed up optimization. The run with the highest multinomial log-likelihood is kept as the best fitting model.

The best-fitting demographic model parameters and the optimal scaling parameter θ_*SYN*_ were savedfor DFE inference. From the θ_*SYN*_ value and the length of synonymous sequence, *L*_*SYN*_, *N*_*anc*_ can becomputed as 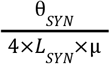 where µ = 1. 5 × 10^−8^. *L*_*SYN*_ is calculated via saved values of exon andintron length from the simulated segments, which are summed over 22 segments to form a single replicate and is found by calculating *L*_*SYN*_ = 0. 302 × *L*_*exon base pairs*_ + *L*_*intron base pairs*_ .

#### Likelihood Ratio Test

We performed a likelihood ratio test (LRT) to test for a significant demographic model fit improvement. We calculated the test statistic *D* = 2(*LL*_3_ − *LL*_2_) where *LL*_*n*_ is the log-likelihood of the best fitting model parameters with epochs, *n. D* asymptotically follows a *X*^2^ distribution [40]. Because the three-epoch model has two additional parameters compared to the two-epoch model, there are 2 degrees of freedom, resulting in a critical value of ∼6 at *p* = 0.05.

### Distribution of fitness effect inference

We fit a gamma-distributed DFE to our nonsynonymous SFS conditional on the two and three epoch demographic fits using fit∂*a*∂*i[8]*. Briefly, fit∂*a*∂*i* precomputes a spectra cache produced from 100 log-spaced selection coefficients (γ = 2*N*_*anc*_ *s*) between 1 × 10^−5^ and 500. We used a Poisson likelihood function in which 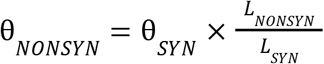 to fit a DFE that best matches the SFS shape and number of SNPs at each frequency of the nonsynonymous SFS. The mutational target ratio, 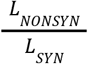, for nonsynonymous SFS scaling was derived from saved values of exon and intron base pair length for each simulation segment where *L*_*SYN*_ = 0. 302 × *L*_*exon base pairs*_ + *L*_*intron base pairs*_ and *L*_*NONSYN*_ = 0. 698 × *L*_*exon base pairs*_.Optimization perturbs the shape and scale parameters within bounds. Shape is constrained between [1 × 10^−3^, 100] and scale between [1 × 10^−2^,1 × 10^6^] in 2*N*_*e*_ |*s*| units.

### Folding the site frequency spectrum

We chose to use folded SFS which summarizes minor allele count from 1 to *n*/2 for our main analyses because a population expansion can model an excess of rare variants but not high frequency variants. Folding was done with ∂*a*∂*i*’s .fold function.

### Comparing simulated and inferred distribution of fitness effects

We were interested in whether the model could capture the true simulated DFE proportions in the presence of selective sweeps. We divided the fit∂*a*∂*i* inferred DFE scale parameter by 2*N*_*anc*_ torecover the unscaled parameter where *N*_*anc*_ = θ_*SYN*_ / 4*L*_*SYN*_ µ. ∂*a*∂*i* parameterizes genotype fitnessesas 1, 1 + 2*hs*, 1 + 2*s*, whereas SLiM uses 1, 1 + *hs*, 1 + *s*. We multiplied the inferred fit∂*a*∂*i* scale by 2 to convert selection coefficients to SLiM units.

We binned the resulting gamma distribution parameters into probability masses within neutral (−10^−5^ < *s* ≤ 0),− 10^−4^ < *s* ≤ − 10^−5^, − 10^−4^ < *s* ≤ − 10^−3^,− 10^−3^ < *s* ≤ − 10^−2^, strongly deleterious (*s* ≤ − 10^−2^). We compared the model bins to those of the true simulated DFE.

### Detecting SFS skews towards rare variation

Linked selection distorts SFS towards rare frequency alleles. To compare skews across simulation regimes, we followed the approach done by Daigle and Johri (2025) [41] by calculating the summary statistic 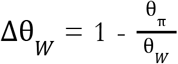. This value has been shown to be less sensitive to the number of SNPs than Tajima’s D [42]. θ_π_ is the nucleotide diversity[43] and θ_*W*_ is Watterson’s θ[44].

## Results

### Population expansion model captures overabundance of rare frequency alleles in single sweep simulations

We first asked whether a misspecified demographic model would absorb the SFS distortions produced by a single completed sweep. All simulations were run under a constant population size, and we fit one-size-change (two-epoch) and two-size-change (three-epoch) models. Because both models include constant size as a special case (ν = 1), any inferred departure from ν = 1 reflects linked selection being absorbed into demography.

As a control, we inserted a neutral mutation (*s*_*ben*_ = 0) in the center of a segment containing neutraland deleterious variation drawn from a known DFE [8]. These simulations included background selection (BGS) but no sweep. BGS slightly shifted the SFS towards rare variation, more strongly within the lower recombination rate simulations. The median Δθ _*W*_ was 0.004 in the *r* = 1 × 10^−8^ simulations and 0.028 in *r* = 1 × 10^−9^ simulations (Figure 1A; Table S2). The two-epoch model absorbed this shift as a slight expansion: *median* ν = 1. 19 at *r* = 1 × 10^−9^ and *median* ν = 1. 09 at *r* = 1 × 10^−8^ (Figure 1B; Table S4). The two-epoch SFS fit the simulated spectra closely, with median Pearson residuals between −0.32 and 0.25 in every frequency bin (Figure 1C; Figure S3; Table S5). The three-epoch model was preferred by a likelihood-ratio test in 2/20 replicates at *r* = 1 × 10^−8^ and 0/20 at *r* = 1 × 10^−9^ (Figure S4; Table S6). The two-epoch model fit the data better than a one-epoch model in some replicates, 2/20 at *r* = 1 × 10^−8^ and 9/20 at *r* = 1 × 10^−9^ (Figure S20; Table S8).

**Figure 1.**
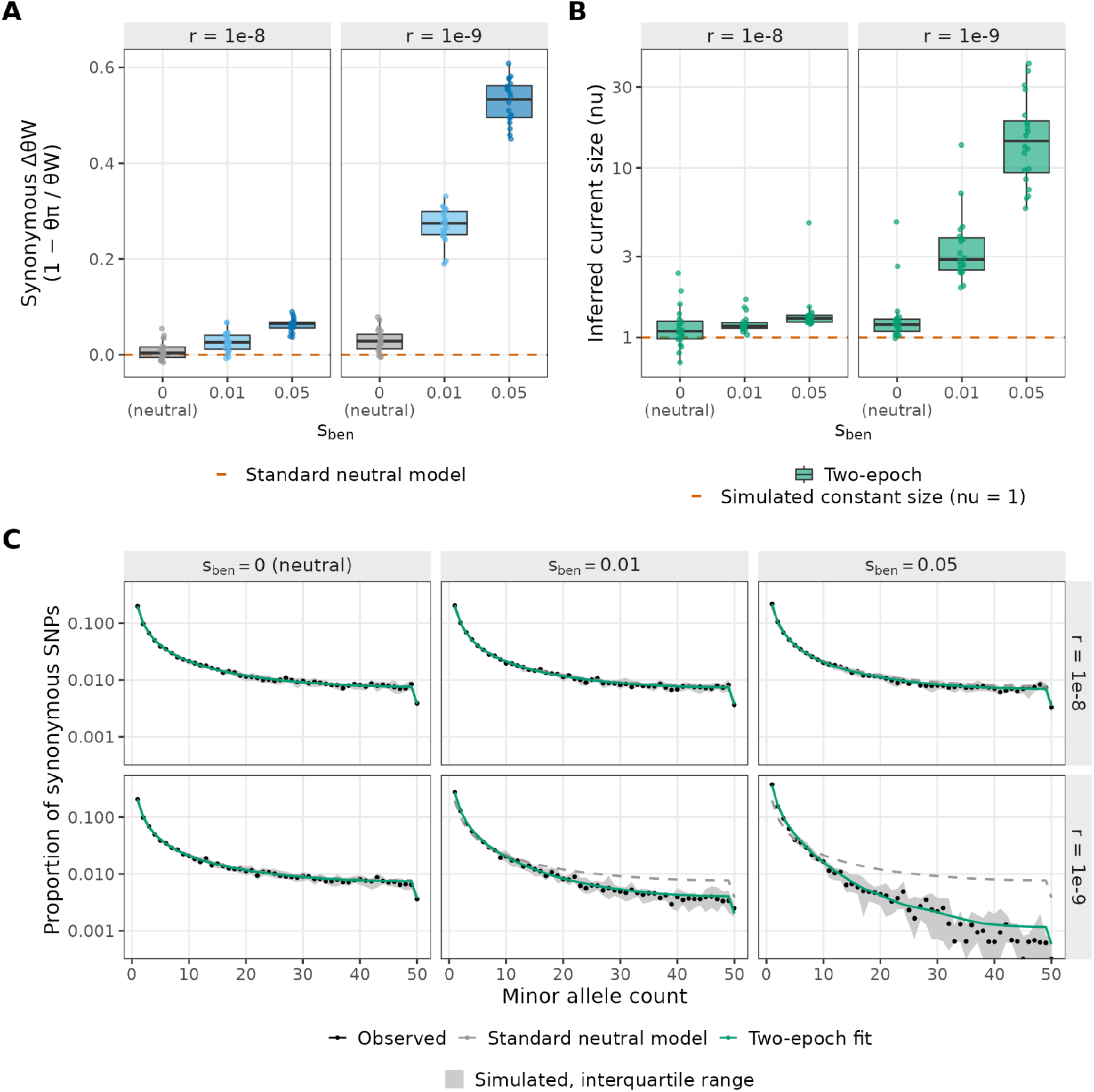
Two-epoch demographic model infers a population expansion after a single selective sweep. Two-epoch demographic inference from simulations of a constant size population (*N*=10,000 diploids). A single beneficial mutation (or neutral mutation in the controls) was placed in the center of the segments after the burn-in period. We sampled at fixation of the beneficial mutation; neutral controls were sampled 2,400 generations after the mutation was introduced. All statistics use the folded synonymous SFS (exonic synonymous and intronic sites) from *n* = 100 genomes, within ±1Mb of the center of the simulation. There are 20 replicates per condition. **(A)** The excess of rare synonymous variants, Δθ_*W*_, by beneficial mutation selection coefficient,*s*_*ben*_, andrecombination rate, *r*. **(B)** Current inferred population size (ν) relative to the ancestral population size inferred by fitting a two-epoch model to the synonymous SFS with ∂*a*∂*i*, reported in log-scale. **(C)** Folded synonymous SFS, shown as the proportion of synonymous SNPs at each minor allele count (log scale). Columns are sweep *s* and rows are recombination rate *r*. Points show the median observed proportion across replicates, and shading shows the interquartile range (IQR) of simulated spectra. The green line is the two-epoch fit and grey dashed line expected SFS under the standard neutral model. Both are the median across replicates. In A and B, each point is a replicate. Boxes show the median and IQR and whiskers reach 1.5x the IQR.

We next inferred the demography in simulations with a single completed selective sweep of increasing strength, *s*_*ben*_ = {0. 01, 0. 05}. The selective sweep caused a greater excess of rare alleles than BGS alone (Figure 1A, C). At *r* = 1 × 10^−8^, the median excess was small (Δθ_*W*_ = 0.026 at *s*_*ben*_ = 0. 01, 0.065 at*s*_*ben*_ = 0. 05) and the demographic model absorbed thesedistortions as slight population expansions which increased with stronger selection strength (medianν = 1. 17 at *s* = 0. 01, median ν = 1. 30 at at *s*_*ben*_ = 0. 05). In the lower recombination ratesimulations,*r* = 1 × 10^−9^, the distortions were much larger, with a medianΔθ_*W*_ of 0.274 (*s*_*ben*_ = 0. 01) and 0.533 (*s*_*ben*_ = 0. 05). The inferred current size was 2.9x (*s*_*ben*_ = 0. 01) and14.5x (*s*_*ben*_ = 0. 05) the ancestral size (Figure 1B; Table S4). The two-epoch fit was close to thesimulated SFS (Figure 1C), so the skew toward low-frequency variants from the sweep was absorbedalmost entirely byν. When *s*_*ben*_ = 0. 05, there were small overestimations of the proportion ofalleles at frequencies of 0-1% (median Pearson residual +1.42) and 20-35% (+1.46) and underestimation of those at frequency 1-5% (−0.78, −1.57) and 35-50% (−1.53) (Figure S3). Adding an additional size change (three epoch) improved model fit in 7/20 replicates at *s* = 0. 05 and1/20 at *s*_*ben*_ = 0. 01. Estimates of ν were similar (11.7 and 2.85) but failed to coverage in 8/20replicates (Figure S4; Tables S4 and S6).

We also evaluated how the window and sample size affected demographic inference. When we increased the window to ± 3 Mb or ± 5 Mb from the center of the sweep, we added synonymous sites that were less linked to the sweep. As expected, this diluted the SFS distortion and decreased the inferred population expansion (Figure S5B; Table S4). Demographic inferences with a larger sample size were not substantially different from those with a sample size of 100. Increasing the sample size to *n* = 1000 yielded similar estimates to *n* = 100 (Figure S5A; Table S4).

We also inferred demography from the unfolded SFS, which summarizes derived allele count from 1 to *n* - 1 rather than minor allele counts from 1 to *n*/2. In the neutral controls, the two-epoch model fit the unfolded synonymous SFS closely (Figure S12) and yielded population size estimates similar to those from the folded SFS (unfolded, ν = 1.10 folded, ν = 1.09 at *r* = 1 × 10^−8^ ; unfolded, ν = 1.22 folded, ν = 1.19 at *r* = 1 × 10^−9^; Figure S5C; Table S4). After a sweep, the SFS was U-shaped, with an excess of both low and high frequency derived alleles (Figure S12). The two-epoch model captured the excess of rare derived alleles but not the excess of high frequency derived alleles, which an expansion predicts should be depleted (Figure S12). As a result, demographic inference absorbed less of the sweep effects when using the unfolded SFS. At *r* = 1 × 10^−9^, the unfolded SFS gave smaller population expansion estimates than with the folded SFS (folded, ν = 2.88, unfolded ν = 2.05 at *s*_*ben*_ = 0. 01; folded, ν = 14.5 unfolded ν = 4.20 at *s*_*ben*_ = 0. 05). At r = 1 × 10^−^8, folded and unfolded results were similar (folded ν = 1.17, unfolded ν = 1.14 at *s*_*ben*_ = 0. 01 and folded, ν = 1.30, unfolded ν = 1.40 at *s*_*ben*_ = 0. 05; Figure S5C; Table S4).

### The effect of a selective sweep on the inferred distribution of fitness effects

We next used fit∂*a*∂*i* [8] to infer a gamma-distributed DFE for new deleterious mutations from the nonsynonymous SFS of the single sweep simulations, conditional on the best-fitting two-epoch demographic model for each replicate. Selection coefficients of deleterious mutations were simulated from the gamma-distributed human DFE of Kim et al. (2017). We binned the probability mass of each inferred DFE into five classes of *s* and compared these with the true simulated DFE: nearly neutral (− 10^−5^ < *s* ≤ 0) 0.209; − 10^−4^ < *s* ≤ − 10^−5^ 0.111; − 10^−4^ < *s* ≤ − 10^−3^ 0.170; − 10^−3^ < *s* ≤ − 10^−2^ 0.248; strongly deleterious (*s* ≤ − 10^−2^) 0.262. Deviations from thesimulated DFE beyond the no-sweep controls indicate biases related to genetic hitchhiking.

In the control simulations containing only neutral and deleterious variation, the inferred DFE recovered the simulated DFE. At *r* = 1 × 10^−8^, every bin was within 0.01 of the true proportion. At *r* = 1 × 10^−9^, consistent with stronger BGS, the inferred DFE slightly underestimated the proportion of moderately deleterious mutations (− 10^−3^ < *s* ≤ − 10^−4^: inferred = 0.158, true = 0.170; − 10^−2^ < *s* ≤ − 10^−3^: inferred = 0.232, true = 0.248) and overestimated the proportion of strongly deleterious mutations (inferred = 0.279, true = 0.262), but the true proportions were within the interquartile range (IQR) values (Figure 2A, Table S7). The median fitted shape and −E[s] parameters were also close to the simulated values (inferred shape = 0.182 when *r* = 1 × 10^−8^, inferred shape = 0.183 when *r* = 1 × 10^−8^, true shape = 0.186; inferred −E[s] = 0.0122, 0.0165 true = 0.0131; Figure 2B).

**Figure 2.**
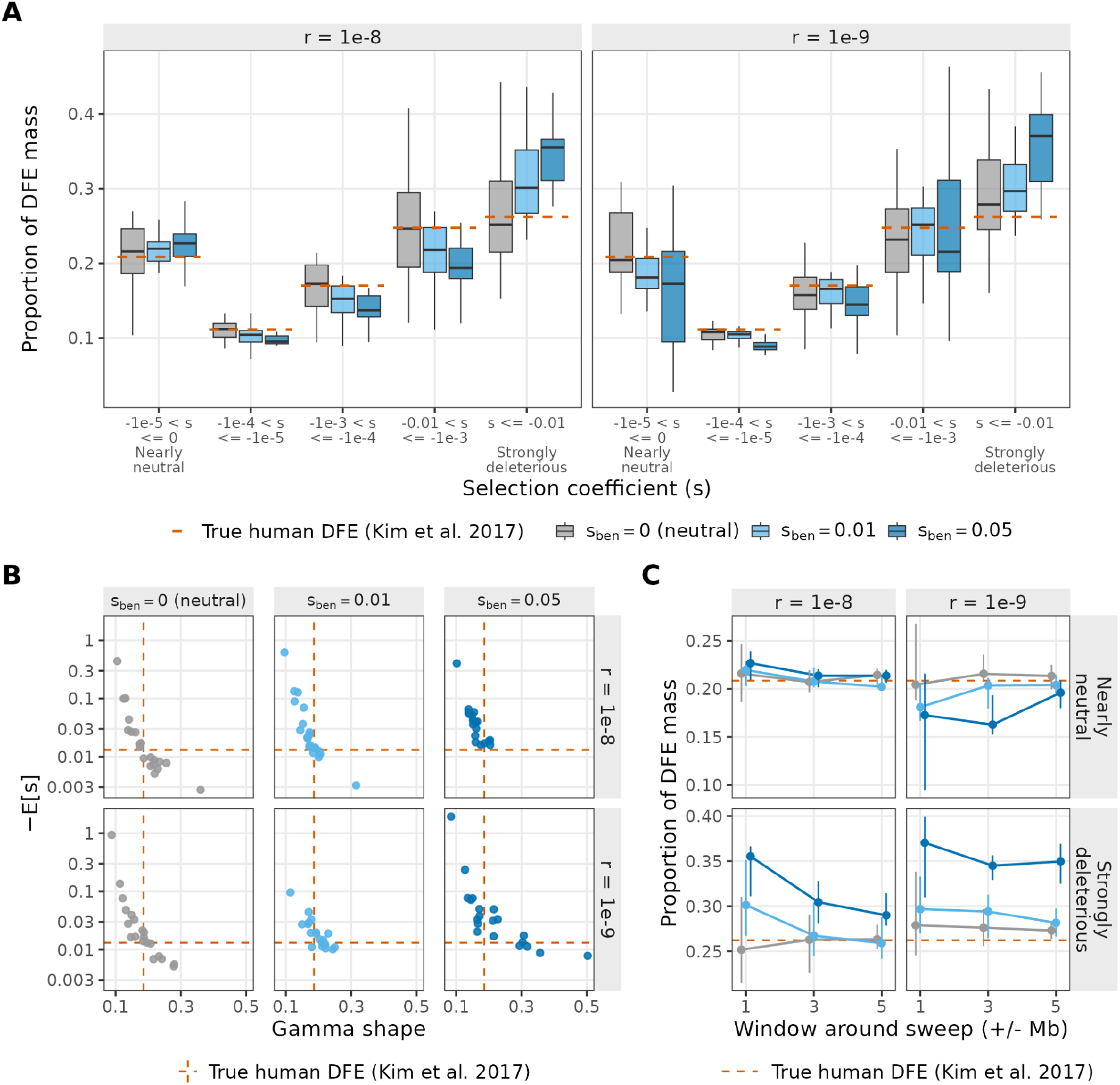
The influence of a completed sweep on the inferred DFE. Gamma distributed DFE inferences on the nonsynonymous SFS, conditioned on the two-epoch demographic models from Figure 1. Selection coefficients are in SLiM (1+*hs*) units. Dashed orange lines represent the true simulated human DFE (Kim et al. 2017); gamma shape = 0.186, E[s] = -0.0131. Panels A and B use SNPs within ±1 Mb of the segment center. Beneficial mutation selection coefficient, *s*_*ben*_, is shown by color in A and C. (**A**) Proportional DFE in five selection-coefficient bins, from nearly neutral to strongly deleterious. Panels are recombination rates. Boxes show the median and IQR across replicates, and whiskers reach 1.5x the IQR. (**B**) Inferred gamma shape plotted against the inferred *-E[s]* (log-scale). Each point is one replicate. Columns are *s*_*ben*_ and rows are recombination rate, *r*. (**C**) Proportion of DFE mass in the nearlyneutral (top row) and strongly deleterious (bottom row), against the width of the window around the sweep (±1,3,5 Mb). Points show the median across replicates and bars the IQR. Points are offset horizontally.

Next, we examined the inferred DFEs in the presence of a single completed selective sweep. Broadly speaking, the inferred DFEs are mostly near the true value used to simulate the data, indicating that the presence of un-modeled selective sweeps does not massively bias inference (Figure 2A). Consistent with this pattern, estimates of the shape and -E[s] values fall near the true values (Figure 2B). However, we note a slight shift in the inferred DFEs toward more strongly deleterious mutations. When *s*_*ben*_ = 0. 01, this skew is modest: the strongly deleterious inferred proportion was 0.301 (IQR: 0.267-0.351) at *r* = 1 × 10^−8^ and 0.370 (0.310-0.399) at *r* = 1 × 10^−9^, with the true value, 0.262, just below the IQR. When the beneficial mutation was more strongly selected (*s* _*ben*_ = 0. 05, above the *E*[|*s*|] of the simulated DFE, 0.013), the overestimation was larger (inferred proportion = 0.355 (IQR: 0.311-0.366) at *r* = 1 × 10^−8^ and inferred proportion = 0.370 (IQR: 0.310-0.399) at *r* = 1 × 10^−9^, with less mass in− 10^−3^ < *s* ≤ − 10^−4^ (inferred proportion = 0.137 at *r* = 1 × 10^−8^ and inferred proportion = 0.145 at *r* = 1 × 10^−9^, true proportion = 0.170). These estimates had a larger −E[s] than the true value simulated (median inferred = 0.040 at *r* = 1 × 10, median inferred = 0.032 at *r* = 1 × 10^−9^, true = 0.013; Figure 2B). The inferred proportion in the nearly neutral bin depended on the recombination rate. It was slightly overestimated at *r* = 1 × 10^−8^ (inferred proportion = 0.220 and inferred proportion = 0.227 for *s*_*ben*_ = 0. 01 and 0.05, true proportion = 0.209) and underestimated at *r* = 1 × 10^−9^ (inferred proportion = 0.181 and 0.173) (Figure 2A). These biases became less pronounced when including sites farther from the sweep, particularly at *r* = 1 × 10^−8^ : for *s*_*ben*_ = 0. 05 the strongly deleterious proportion (true = 0.262) fell from 0.355at ±1 Mb to 0.304 at ±3 Mb and 0.290 at ±5 Mb, compared with 0.370, 0.345 and 0.350 at*r* = 1 × 10^−9^(Figure 2C; Figure S7; Table S7). This suggests that the proportion of variants linked to the sweeping beneficial variant affects the extent of the bias in inference of the deleterious DFE.

A three-epoch demographic model gave similar DFEs with similar IQRs compared to the two-epoch demographic model (strongly deleterious inferred proportion = 0.354 at *r* = 1 × 10^−8^ and 0.385 at *r* = 1 × 10^−9^ for *s*_*ben*_ = 0. 05; Figure S14). With a larger sample (*n* = 1000) the IQRs narrowed (e.g. the strongly deleterious IQR width in the *r* = 1 × 10^−9^ controls fell from 0.093 to 0.051). The overestimation of the proportion of strongly deleterious mutations persisted but was smaller (inferred = 0.307 at *r* = 1 × 10^−8^and 0.334 at *r* = 1 × 10^−9^ for *s*_*ben*_ = 0. 05), whereas the underestimation of the proportion of nearly neutral mutations at *r* = 1 × 10^−9^ was larger (inferred = 0.164 and 0.113 for s_ben = 0.01 and 0.05) (Figure S8; Table S7).

Thus far, our analyses considered the folded SFS. Next, we examined the impact of a single sweep on inference when using the unfolded SFS. After a sweep, inferences of the DFE using the unfolded SFS were more biased than when using the folded SFS (Figure S5C), reflecting the demographic model not capturing the sweep SFS distortions (Figure S12). For *s*_*ben*_ = 0. 05, weakly tomoderately deleterious bins were underestimated and shifted towards the strongly deleterious bins where the proportion was 0.374 unfolded and 0.355 folded (true proportion = 0.262). Median −E[s] using the unfolded SFS was inferred to be 0.063 at *r* = 1 × 10^−8^ and 8.8 at *r* = 1 × 10^−9^(true= 0.013; Figure S11B).

### Distribution of fitness effect inference remains robust with recurrent beneficial mutations

After evaluating DFE inference biases caused by a single completed selective sweep, we inferred DFEs from simulations with recurrent beneficial mutations. A proportion, *p*_*ben*_, of newnonsynonymous mutations had a fixed beneficial selection coefficient, *s*_*ben*_. We chose five pairs with*p*_*ben*_ × *s*_*ben*_ = 10^−6^, matching levels of human polymorphism and divergence [33], ranging from*s*_*ben*_ = 0.00032 (*p*_*ben*_ = 0.003125) to *s*_*ben*_ = 0.05 (*p*_*ben*_ = 0.00002). More strongly selected beneficial mutations arose less often but were more likely to fix and fixed faster: at *r* = 1 × 10^−8^, fixations rose from 1010 to 1323 across 20 replicates, and the median time from origin to fixation fell from 23,360 to 600 generations (Figure S15; Table S9). We inferred demography from the synonymous SFS composed of 22 10 Mb segments and the DFE conditional on the two-epoch demography, and compared both with controls at the same window size and generation runtime.

Recurrent sweeps shifted the synonymous SFS slightly toward rare variants (Figure S1; Figure S16A). At *r* = 1 × 10^−8^, Δθ_*W*_ rose from 0.005 (IQR 0.002–0.011) in the control simulations to 0.021 (0.011–0.026) at*s*_*ben*_ = 0. 05, and only at *s*_*ben*_ = 0. 05 did the IQR not overlap that of the controlsimulations. At*r* = 1 × 10^−9^,Δθ_*W*_ rose from 0.028 (0.020–0.031) in the controls to 0.044(0.034–0.050), 0.055 (0.044–0.067) and 0.090 (0.077–0.104) at*s* = 0. 0032, 0.01, 0.05, withIQRs above the controls (Table S2). The two-epoch model absorbed these shifts as slight population expansions. At *r* = 1 × 10^−8^, the inferred current size was 1.03- (*s*_*ben*_ = 0. 00032) to 1.09-fold (*s*_*ben*_ = 0. 05) the ancestral size, close to that inferred in the control simulations (1.03). At *r* = 1 × 10^−9^, it was 1.14- to 1.37-fold, compared with 1.12 in the control simulations (Figure 3A; Table S4).

**Figure 3.**
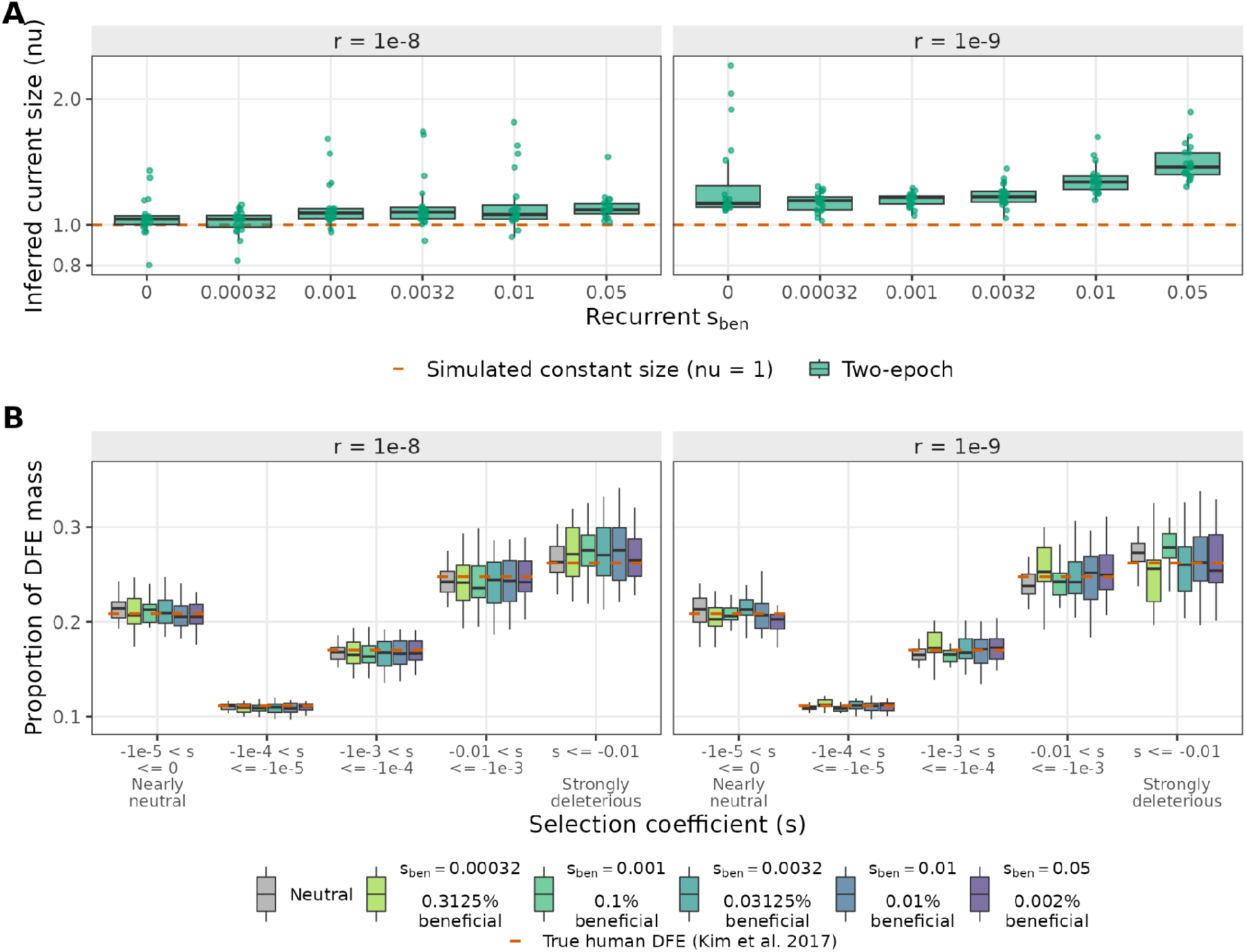
Recurrent sweeps occurring at a human-like rate have little effect on the inferred demography or DFE. Results come from simulations of a constant-size population (*N*=10,000 diploids). Each replicate contains deleterious mutations from the human DFE (Kim et al. 2017), with beneficial mutations of strength arising throughout the simulation. In each condition, a fraction, *p*, of new nonsynonymous mutations is beneficial with selection coefficient, *s*_*ben*_, where *log*_10_ (*s* × *p*) = − 6. Samples were taken at generation 102,400. The neutral replicates are thesame as in Figure 1, but with a window of size ± 5 Mb from the segment center. All statistics use the folded SFS from *n* = 100 haploid genomes over the full 10 Mb segment. Each replicate contains 22 segments. There are 20 replicates per condition. **(A)** Current inferred population size (ν) relative to the ancestral population size inferred by fitting a two-epoch model to the synonymous SFS with ∂*a*∂*i*, reported in log-scale. **(B)** Proportional DFE in five selection-coefficient bins, from nearly neutral to strongly deleterious. The DFE was fitted to nonsynonymous SFS with fit∂*a*∂*i*, conditioned on the two-epoch demography in A, with selection coefficients in SLiM units. Color gives the recurrent-sweep condition. Dashed orange lines mark the true deleterious human DFE (Kim et al. 2017). Panels are recombination rates. In both panels, boxes show the median and IQR across replicates, and whiskers reach 1.5x the IQR.

The inferred DFEs were close to the simulated DFE in every condition (Figure 3B; Table S7). At *r* = 1 × 10^−8^, the true simulated proportion was within the IQR of the inferred proportions in every bin of the DFE. At *r* = 1 × 10^−9^, the no-sweep controls placed the simulated proportion just outside the IQR in two bins, underestimating −10^−4^ < s ≤ −10^−5^ (0.110 [0.1078–0.1111] vs 0.1115) and overestimating the proportion of strongly deleterious mutations (0.273 [0.264–0.283] vs 0.262). The recurrent-sweep DFEs deviated by similar amounts. At *s*_*ben*_ = 0.001 the same two bins and −10^−3^ < s ≤ −10^−4^ did not overlap the simulated value (strongly deleterious 0.278 [0.267–0.293]), and at *s*_*ben*_ = 0.05 the nearly neutral bin was slightly underestimated (0.203 [0.192–0.208] vs 0.209). These deviations did not increase with *s*_*ben*_. The inferred DFEs remained close to the simulated DFEs when using an unfolded SFS, larger sample size (*n*=1000), and three-epoch demographic model as well (Figures S17-S19).

## Discussion

Here we evaluated how selective sweeps affect inference of the deleterious DFE with fit∂*a*∂*i [8]*. We found that DFE inferences were relatively robust to the presence of selective sweeps across a variety of parameter conditions. While the selective sweeps caused distortions to the SFS, fitting a demographic model to linked neural variants largely accounted for much of this distortion. However, closer examination showed slight biases in the inference under certain conditions. For example, when SFS were sampled immediately after fixation of a single beneficial mutation, linked positive selection biased fit∂*a*∂*i* towards overestimating the proportion of strongly deleterious mutations, particularly at lower recombination rates. However, this bias largely disappeared under a more realistic model of recurrent sweeps, with beneficial mutations occurring at a rate expected from human polymorphism and divergence data [33]. fit∂*a*∂*i* remained robust whether beneficial mutations were weakly selected and comprised a large fraction of new mutations, or strongly selected at a low fraction.

The DFE biases present in the single-sweep simulations were consistent with those previously reported for DFE-alpha and polyDFE [10,31,32]. However, our single-sweep approach differed from these studies. Messer and Petrov (2013) and Tataru et al. (2017) specified beneficial alleles occurring as a proportion of nonsynonymous mutations. In contrast, beneficial mutations in our single sweep simulations were conditional-on-fixation and did not contribute to the SFS. The distortions we observed were caused by hitchhiking at linked sites, rather than the combination of hitchhiking and beneficial mutations segregating at high frequency.

We also fit a more flexible demographic model (three-epoch) than Messer and Petrov (2013) who considered only a single size change model. A three-epoch model was preferred over a two-epoch model in 7/20 replicates at the most extreme simulation parameters (*s*_*ben*_ = 0.05, *r* =1e-9; Figure S4, Table S6). However, the three-epoch model was more difficult to fit and produced DFE biases similar to those of the two-epoch model (Figure S14; Table S6, S7). In general, the demographic models fit the folded synonymous SFS well (Figures 2C, S2, S3; Table S5) and did not reach our optimization bounds, yet the small DFE biases persisted (Figure 2; Table S7). This suggests that hitchhiking distorts the SFS at selected sites in ways a demographic model fit to synonymous sites cannot fully capture, so demographic correction alone is insufficient to completely account for sweeps.

The directionality of the DFE bias, a depletion of weakly deleterious mutations and excess of strongly deleterious mutations, can be understood by considering how hitchhiking differs across selective strength and how the fitted demographic model absorbs these differences. Neutral variants linked to the sweep can hitchhike to high frequency, limited only by recombination and random drift [20]. Deleterious variants behave differently. Some deleterious variants will hitchhike, but purifying selection may also counteract positive selection [25,26]. Meanwhile, the excess of rare synonymous variants left by the sweep is fit as a population expansion [17]. Under this expansion, the demographic model expects a high proportion of rare nonsynonymous variants, so rare variants that would indicate weak purifying selection are attributed to demography [21]. In effect, the demographic expansion “overcorrects” for the sweep. The resulting deficit of observed nonsynonymous variants relative to the expansion expectation is best explained by strongly deleterious or lethal mutations, which do not segregate in the sample. Mass is therefore shifted out of the weakly deleterious bins and into the strongly deleterious bins. Similar biases have been reported in selfing populations with pervasive HRI [41] and when including synonymous variants under selection in the demographic inference [45]. Our results add to a general pattern that when synonymous mutations are affected by selection, DFE inference tends to push mass towards the extremes of the distribution.

In contrast to the single-sweep case, DFE inference remained robust under recurrent beneficial mutation occurring at rates consistent with human polymorphism and divergence data [33]. In these simulations, weakly beneficial mutations arose frequently but behaved effectively neutrally. Conversely, strongly beneficial mutations arose less frequently and fixed quickly, so they affected polymorphism only briefly (Table 1; Figure S15; Table S9). In both cases, the inferred demography included a small population expansion, which grew larger as beneficial selection became stronger (Figure 3A; Table S4). However, the DFE was largely unaffected (Figure 3B; Figure S16B; Table S7). This contrasts with the biases reported by Messer and Petrov (2013) and Tataru et al. (2017), likely because their simulations assumed higher rates of adaptation. Because selective sweeps appear to have been rare in recent human history [46], our results suggest that current human DFE inferences are unlikely to be substantially biased by genetic hitchhiking. However, inferences were more variable in regions of low recombination (Figure 2B), where genetic hitchhiking and background selection affect more sites [20,23,25]. Future studies should consider removing regions of low recombination from inference.

Interestingly, we found that inference of the DFE using the unfolded SFS performed worse than when using the folded SFS (Figure S11; Table S7). The unfolded SFS contains more information about high-frequency derived variants, which are especially affected by selective sweeps. Single population demographic models are unable to properly account for these high-frequency derived variants. Consequently, the inferred demographic models do not fit the simulated SFS well (Figure S12). Then, inferences of the DFE for nonsynonymous variants are even more biased (Figure S11; Table S7). The SFS from the best-inferred DFEs also do not match the simulated SFS well either, indicating poor model fit (Figure S13). Thus, our study provides a strong practical recommendation to use the folded SFS when inferring the deleterious portion of the DFE in the potential presence of unmodeled selective sweeps.

Our study has several limitations that suggest future work. First, we considered only a gamma distributed DFE, because it is commonly used in empirical studies [47] and is often preferred even in simulations when the true DFE is more complex [8]. More flexible models, such as a gamma distribution with a neutral or lethal point mass [48], may absorb some of the sweep-induced bias within the point mass component of the distribution. Second, we simulated only a constant population size (*N*=10,000). More complex demographic histories may reduce the efficacy of selection [49] but also make it harder to correct for with a demographic model. Finally, DFE biases in populations with larger ancestral population sizes remain unexplored. Positive selection is more efficient in a larger population. Whether this will reduce the accuracy of fit∂*a*∂*i* is unclear. However, Lin et al. (2026) inferred that mosquitoes, fruit flies, and halictid bees have a lower median deleterious |s| value relative to humans [47]. As they inferred mutations are less deleterious in species with large effective population size, this trend is opposite of the bias we observed in our single sweep simulations, suggesting that sweep-induced bias is unlikely to explain these differences between species.

This study clarifies the accuracy of fit∂*a*∂*i* in the presence of selective sweeps. A single recent sweep may bias the inferred DFE towards strongly deleterious mutations, particularly at low recombination rates and with an unfolded SFS. However, under realistic strengths and proportions of beneficial mutations, DFE inference with fit∂*a*∂*i* remained robust, suggesting that current estimates of the human DFE are unlikely to be substantially biased by linked positive selection.

## Code availability

Scripts are available on Github (github.com/emmaewade/Sweeps_DFE).

## Acknowledgements

The authors thank Abby Bigham and members of the Lohmueller Lab for helpful discussions.

