## Supplementary Figures for "Robustness of the inference of the distribution of fitness effects to selective sweeps"

### Supplementary Information

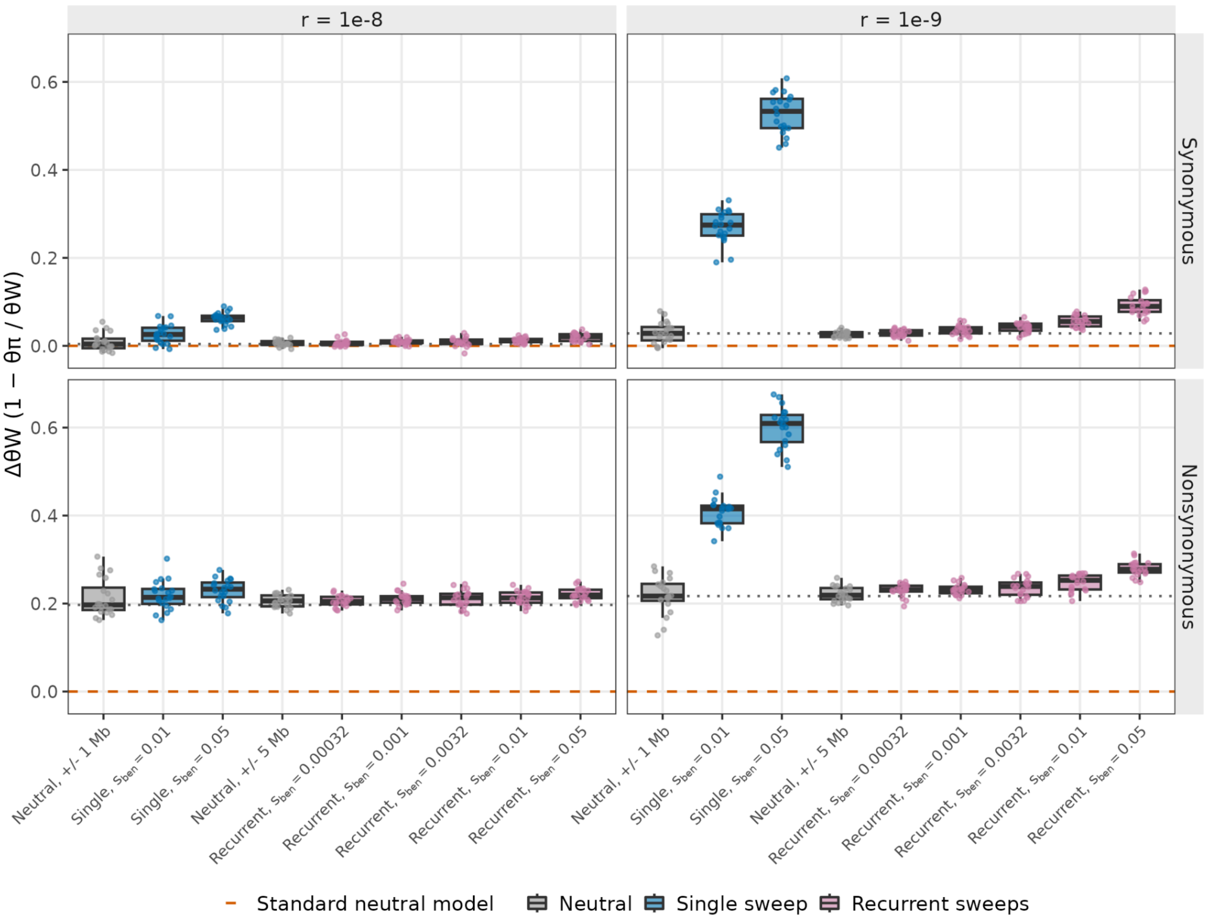

**Figure S1. Selective sweeps produce an excess of rare variation at synonymous and nonsynonymous sites**

Relative excess of rare variants, $\Delta\theta_W$. across selective sweep regimes, computed from the folded SFS ($n$=100) composed of synonymous (top row) and nonsynonymous (bottom row) sites. Columns show the recombination rate. Single-sweep simulations (blue) use sites within +/- 1 Mb from the beneficial mutation. Recurrent-sweep simulations (pink) use the whole 10 Mb (+/- 5 Mb). Neutral controls are shown at both +/- 1 Mb and 5 Mb windows. Boxes show the median and interquartile range across 20 replicates, whiskers extend 1.5x the interquartile range, and points show individual replicates. The dashed orange line marks the standard neutral model expectation. The dotted grey line marks the +/- 1 Mb neutral-control median for each site class and recombination rate.

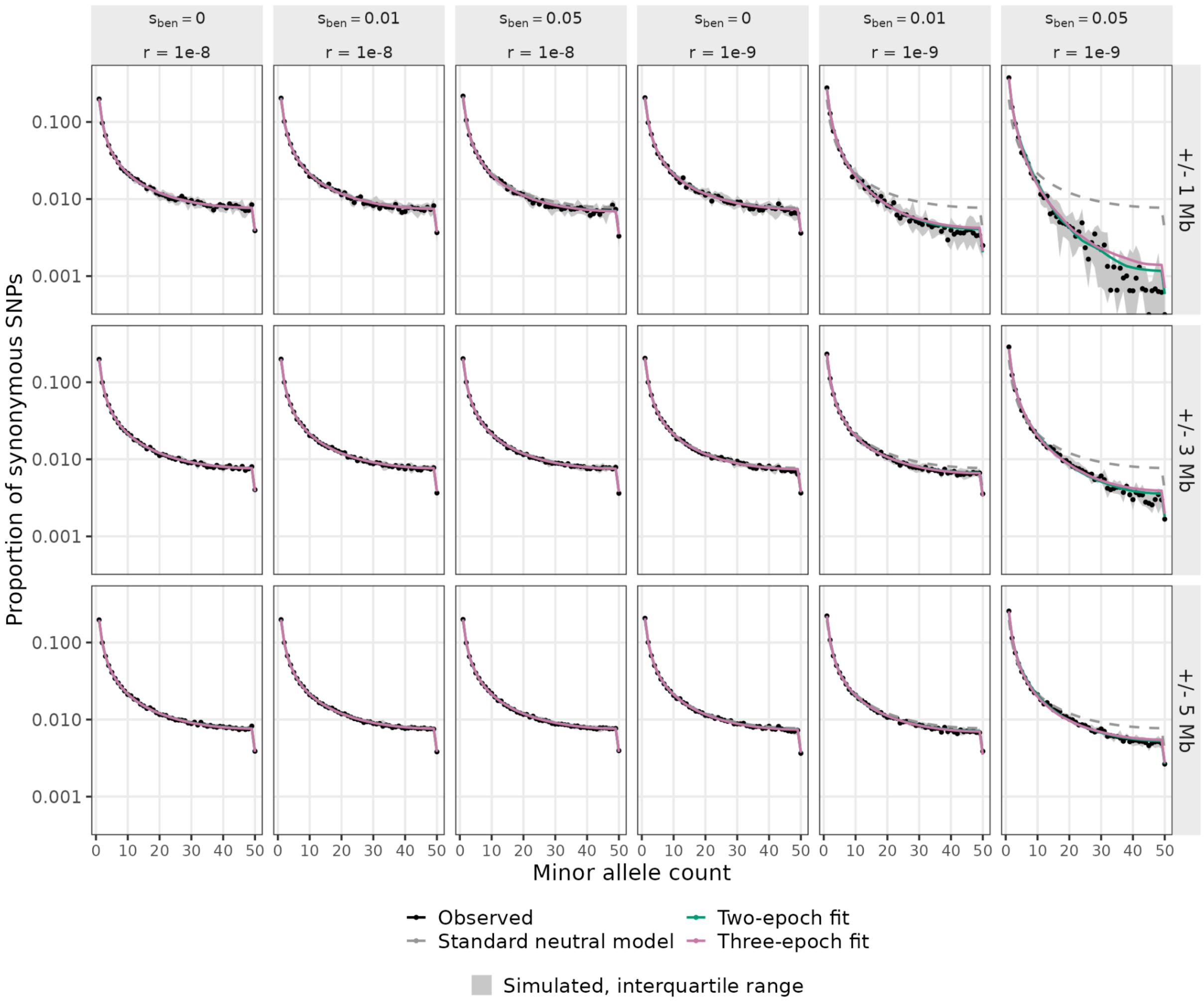

**Figure S2. Two- and three-epoch demographic fits to the synonymous SFS after a single completed sweep**

Folded synonymous SFS ($n$ = 100) from single-sweep simulations, shown as the proportion of synonymous SNPs at each minor allele count (log scale). Columns show the sweep selection coefficient, $s_{ben}$, at each recombination rate, $r$. Rows show the window around the sweep used to build the SFS. Black points show the median observed proportion across 20 replicates, and grey shading shows the interquartile range across replicates. The dashed grey line shows the standard neutral model. Green and pink lines show the median SFS from the two-epoch and three-epoch demographic models fitted to each replicate, respectively.

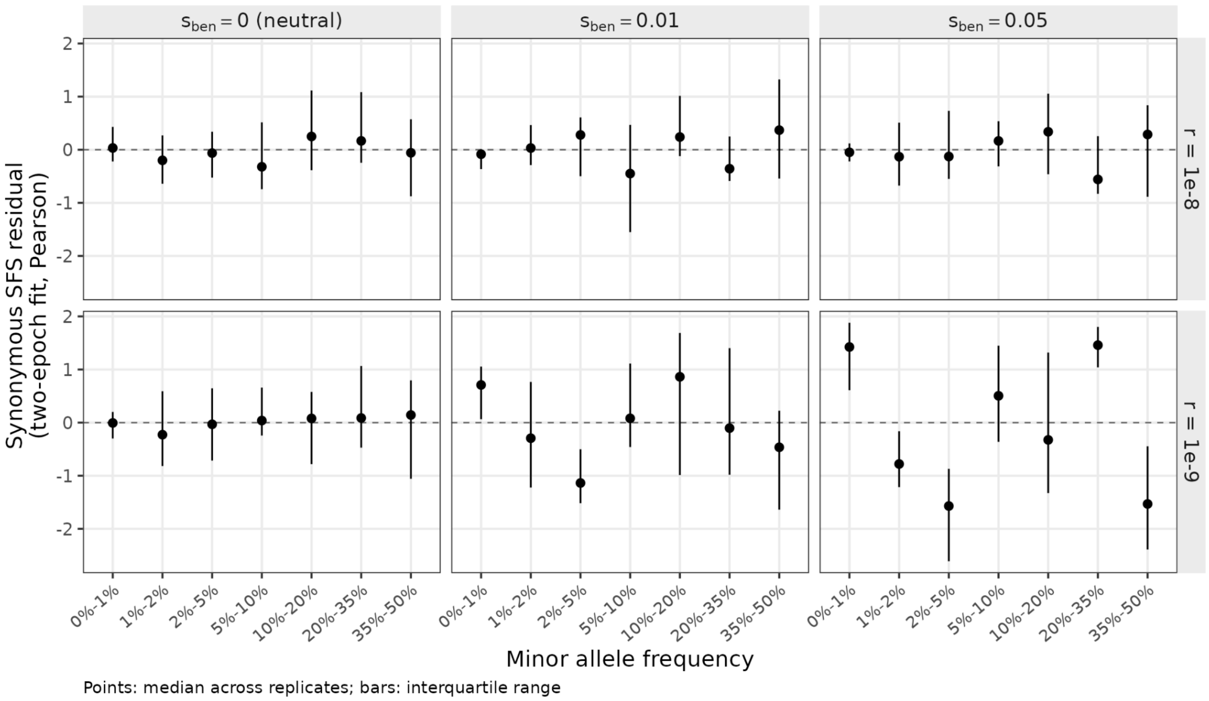

**Figure S3. The two-epoch model fits the synonymous SFS closely**

Pearson residuals of the two-epoch demography fit to the folded synonymous SFS ($n$ = 100) ($\frac{observed-expected}{\sqrt{expected}}$). SNPs are pooled within minor allele frequencies. The first two bins contain singletons and doubletons, respectively. Sites within $\pm$1 Mb of the sweep are included. Columns show the sweep selection coefficient, and rows show the recombination rate. Points show the median across 20 replicates.

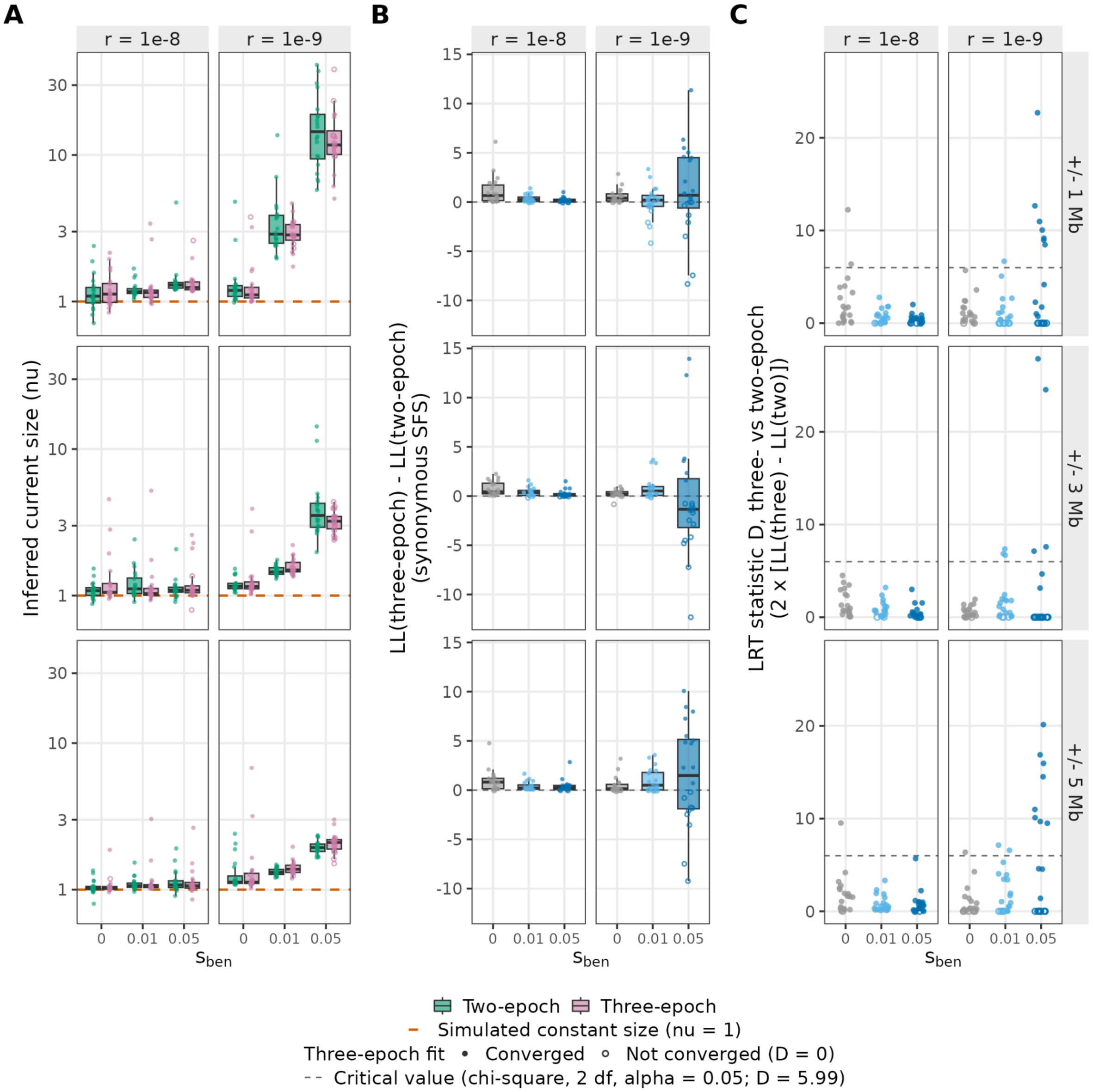

**Figure S4. Two- and three-epoch models infer similar population expansion parameters**

Comparison of two- and three-epoch demographic models fitted to the synonymous SFS ($n$ = 100) after a single completed selective sweep. Columns within panels show the recombination rate. Rows represent the size of the window around the sweep site used to build the SFS. The x-axis show the selection coefficient of the beneficial mutation, $s_{ben}$. **(A)** Inferred current size relative to ancestral, ν, (log-scale) under the two-epoch (green) and three-epoch (pink). The orange dashed line marks the simulated constant size. **(B)** Difference in log-likelihood between the two- and three-epoch fits to the synonymous SFS. The dashed line marks no difference. **(C)** Likelihood-ratio test statistic, $D = 2 \times [LL_3 - LL_2]$. The dashed line marks the $\chi^2$ critical value (2 df, $\alpha = 0.05$, D = 5.99). Open symbols represent replicates that did not converge ( $LL_3 < LL_2$), with D set to 0. In A and B, boxes show the median and interquartile range across 20 replicates, whiskers extend 1.5 x IQR, and points show individual replicates.

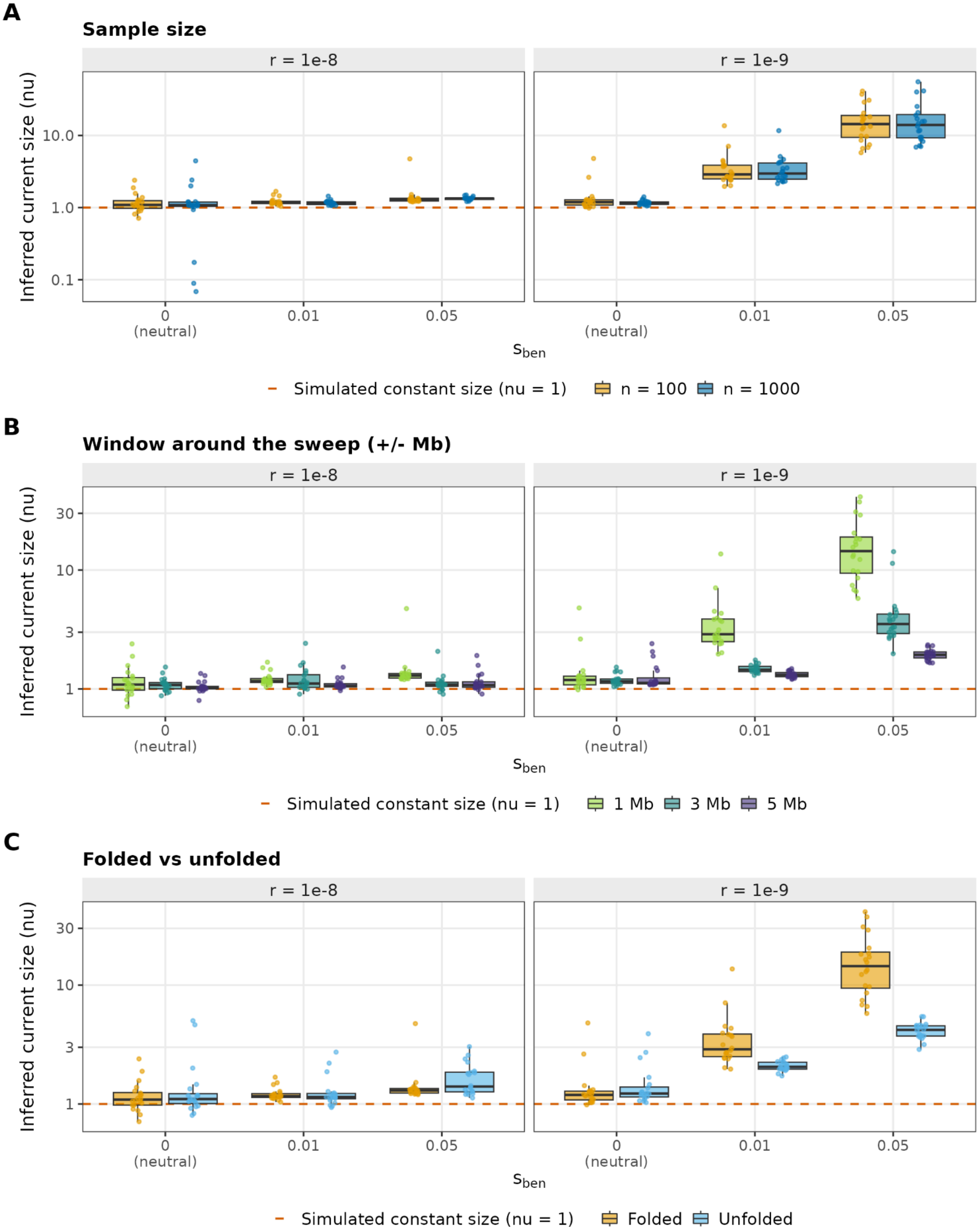
A
Sample size
r = 1e-8
r = 1e-9
Inferred current size (nu)
10.0
1.0
0.1
0
(neutral)
0.01
0.05
sben
Simulated constant size (nu = 1)
n = 100
n = 1000
B
Window around the sweep (+/- Mb)
r = 1e-8
r = 1e-9
Inferred current size (nu)
30
10
3
1
0
(neutral)
0.01
0.05
sben
Simulated constant size (nu = 1)
1 Mb
3 Mb
5 Mb
C
Folded vs unfolded
r = 1e-8
r = 1e-9
Inferred current size (nu)
30
10
3
1
0
(neutral)
0.01
0.05
sben
Simulated constant size (nu = 1)
Folded
Unfolded

**Figure S5. Inferred population expansion depends on window size and folding the SFS, but not on sample size**

Current population size (ν, log scale) inferred by the two-epoch demographic model fitted to the synonymous SFS of single-sweep simulations, under different analysis choices. Columns within panels represent recombination rate, *r*. The x-axis shows the sweep selection coefficient, $s_{ben}$. **(A)** Sample size of *n* = 100 (orange) and *n* = 1000 (blue) with folded SFS and ±1 Mb windows **(B)** Window around the sweep in which SFS are drawn. **(C)** Folded (orange) and unfolded (light blue) SFS with *n* = 100 and ±1 Mb windows. Boxes show the median and interquartile range across 20 replicates, whiskers extend 1.5x the interquartile range, and points show individual replicates. The orange dashed line marks the simulated constant size.

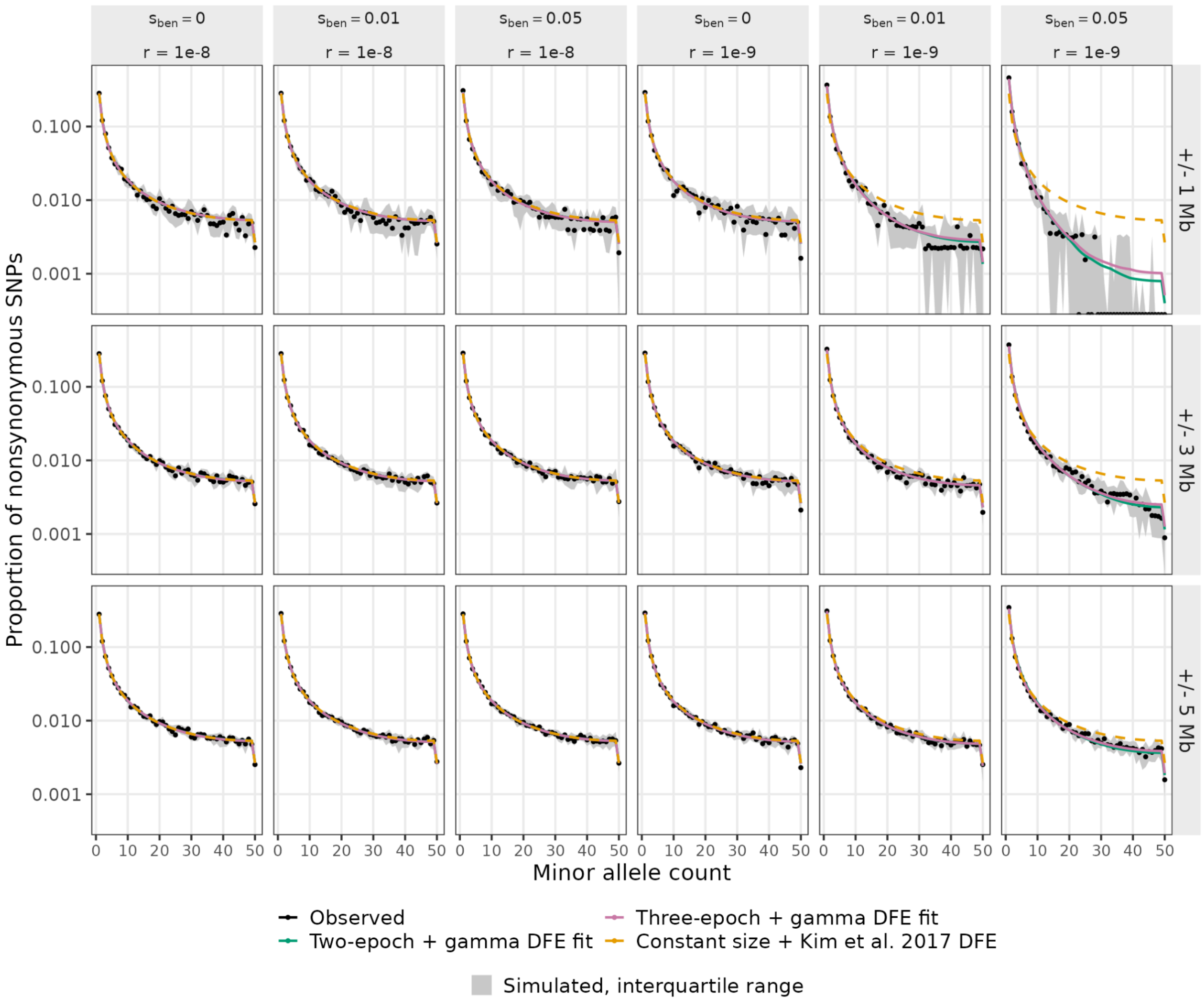

**Figure S6. Gamma DFE fits nonsynonymous SFS after a single completed sweep**

Folded nonsynonymous SFS ($n$ = 100) from single-sweep simulations, shown as the proportion of nonsynonymous SNPs at each minor allele count (log scale). Columns show the sweep selection coefficient at each recombination rate. Rows show the window around the sweep used to build the SFS. Black points show the median observed proportion across 20 replicates, and grey shading shows the interquartile range across replicates. Green and pink lines show the median SFS from gamma DFEs fit conditional to a two-epoch and three-epoch demographic models, respectively. The dashed orange line shows the expected SFS under constant population size with the true human DFE (Kim et al. 2017).

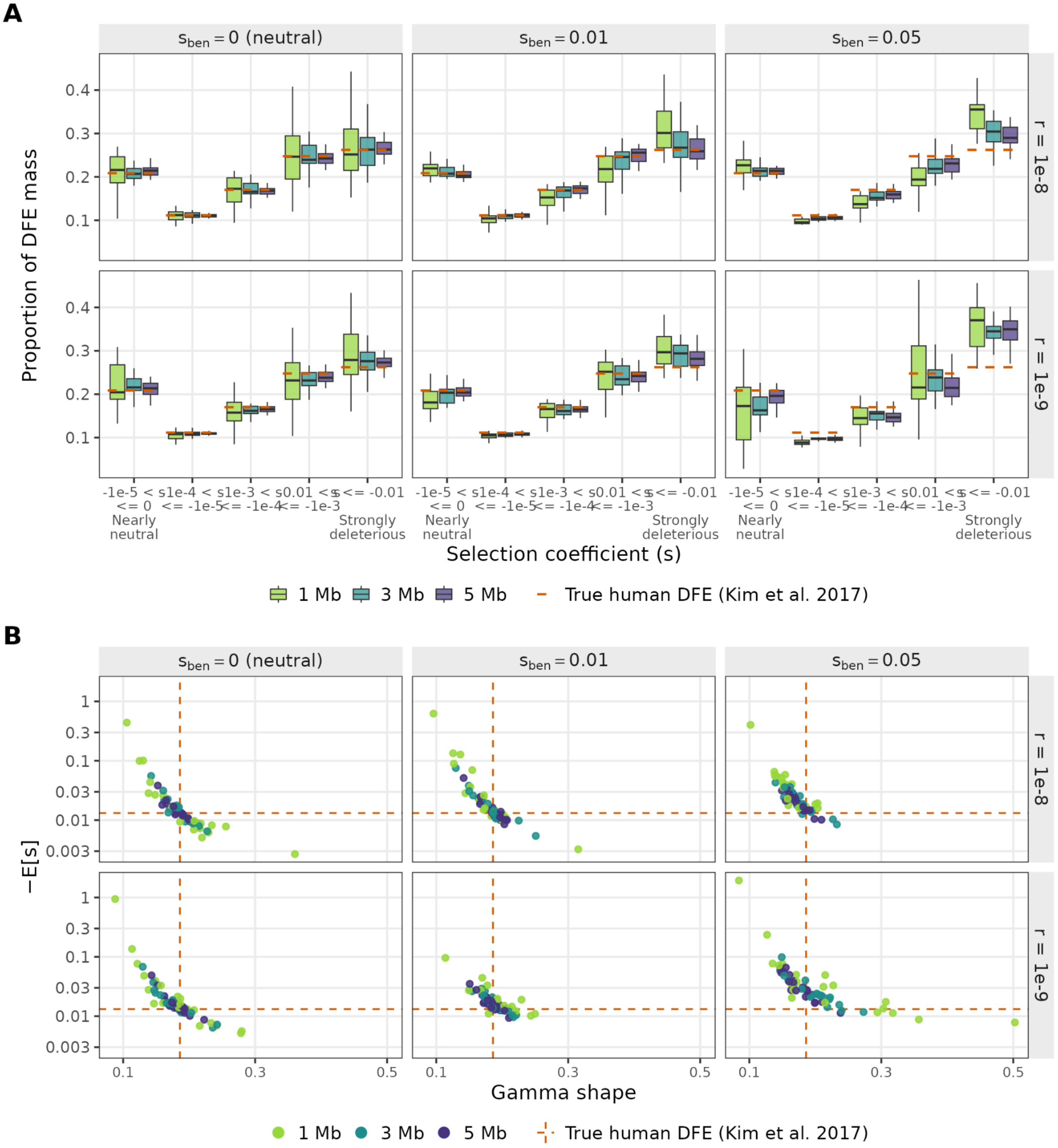

**Figure S7. Inferred DFE bias decreases when sampling sites farther from the sweep**

DFEs inferred after a single completed selective sweep using windows of ±1Mb, ±3Mb, and ±5Mb from the beneficial mutation. Each gamma DFE was fitted to folded nonsynonymous SFS ($n$ = 100), conditional on the two-epoch demography fitted to the synonymous SFS Columns show the beneficial mutation selection coefficient, $s_{ben}$, and rows show the recombination rate, *r*. **(A)** Proportion of DFE mass in each five selection coefficient bins. Boxes show median and interquartile range across 20 replicates, and whiskers extend 1.5x the interquartile range. **(B)** Inferred mean selection coefficient (-$E[s]$, log scale), against gamma shape. Each point represents a replicate. In both panels, orange dashed lines mark the true human DFE (Kim et al. 2017; shape = 0.186, -$E[s]$ = 0.0131).

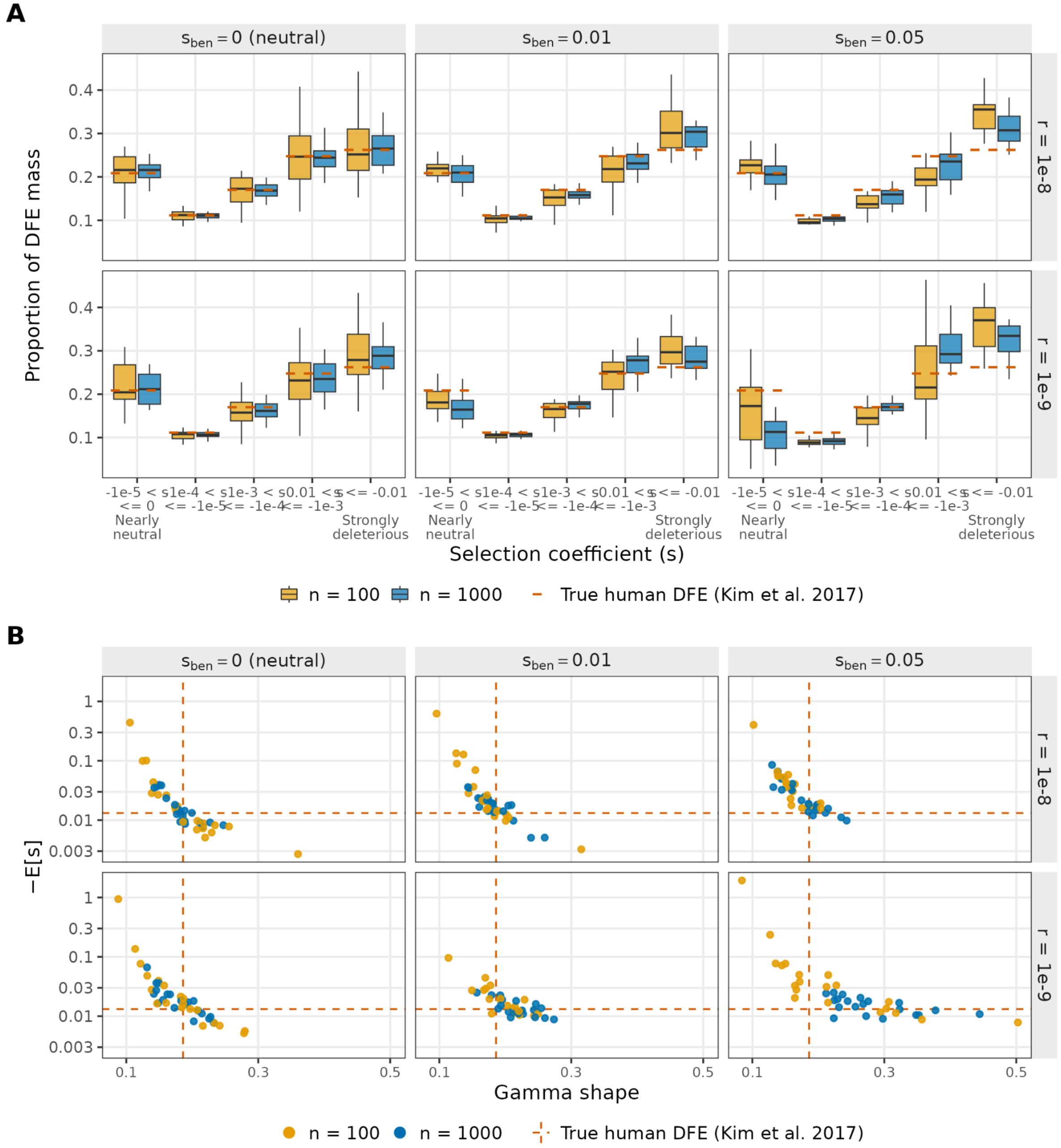

#### Figure S8. Inferred DFEs are similar between sample sizes

DFEs inferred after a single completed selective sweep using sample sizes of $n$ = 100 and $n$ = 1000. Each gamma DFE was fitted to folded nonsynonymous SFS ($\pm\ 1\ Mb$), conditional on the two-epoch demography fitted to the synonymous SFS. Columns show the beneficial mutation selection coefficient, $s_{ben}$, and rows show the recombination rate, $r$. **(A)** Proportion of DFE mass in each five selection coefficient bins. Boxes show median and interquartile range across 20 replicates, and whiskers extend 1.5x the interquartile range. **(B)** Inferred mean selection coefficient (-$E[s]$, log scale), versus the gamma shape parameter. Each point represents a replicate. In both panels, orange dashed lines mark the true human DFE (Kim et al. 2017; shape = 0.186, -$E[s]$ = 0.0131).

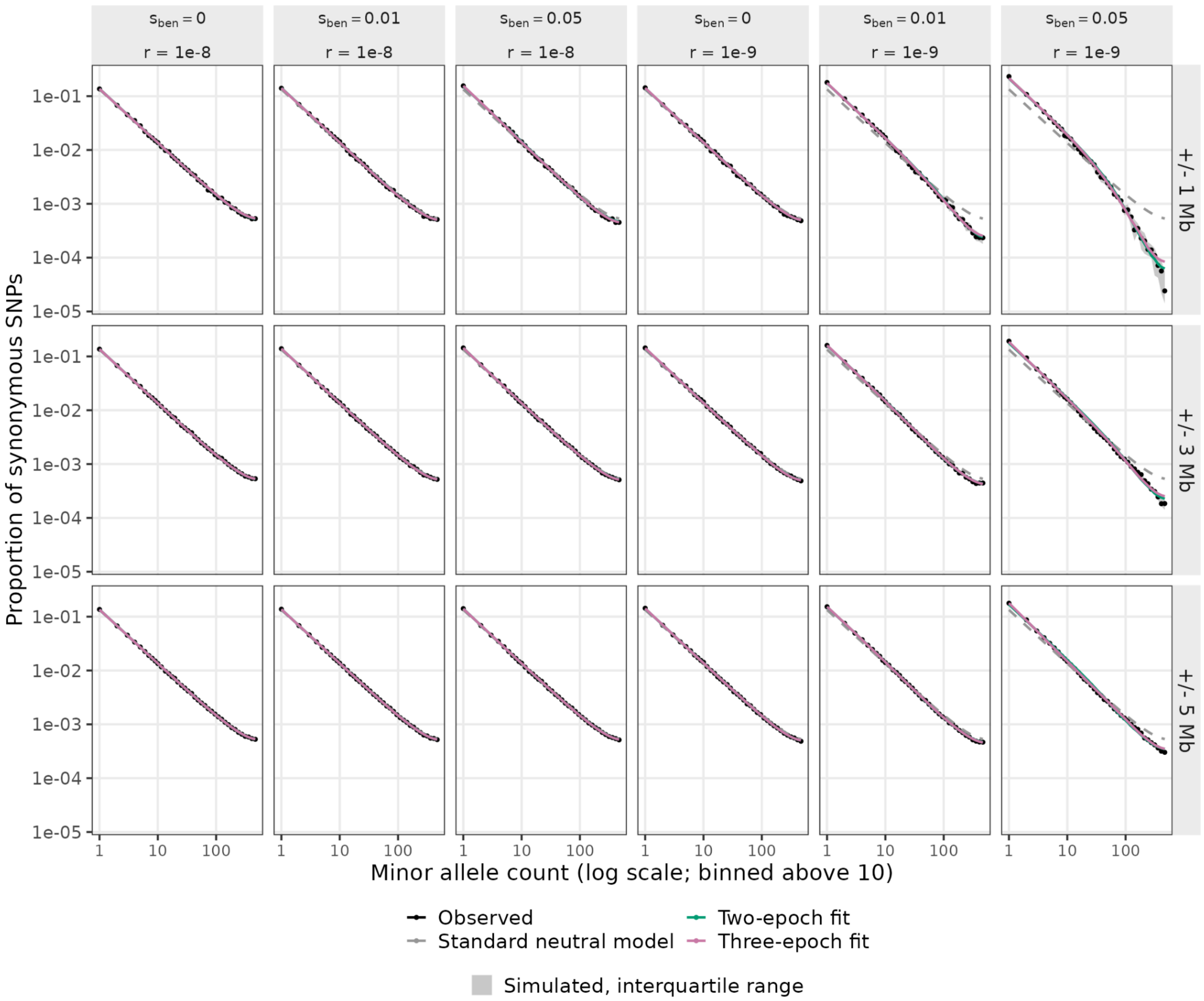

#### Figure S9. Two- and three-epoch demographic fits to the synonymous SFS after a single completed sweep at *n* = 1000

Folded synonymous SFS (*n* = 1000) from single-sweep simulations, shown as the proportion of synonymous SNPs at each minor allele count (log scale). Columns show the sweep selection coefficient at each recombination rate. Rows show the window around the sweep used to build the SFS. Black points show the median observed proportion across 20 replicates, and grey shading shows the interquartile range across replicates. The dashed grey line shows the standard neutral model. Green and pink lines show the median SFS from the two-epoch and three-epoch demographic models fitted to each replicate, respectively.

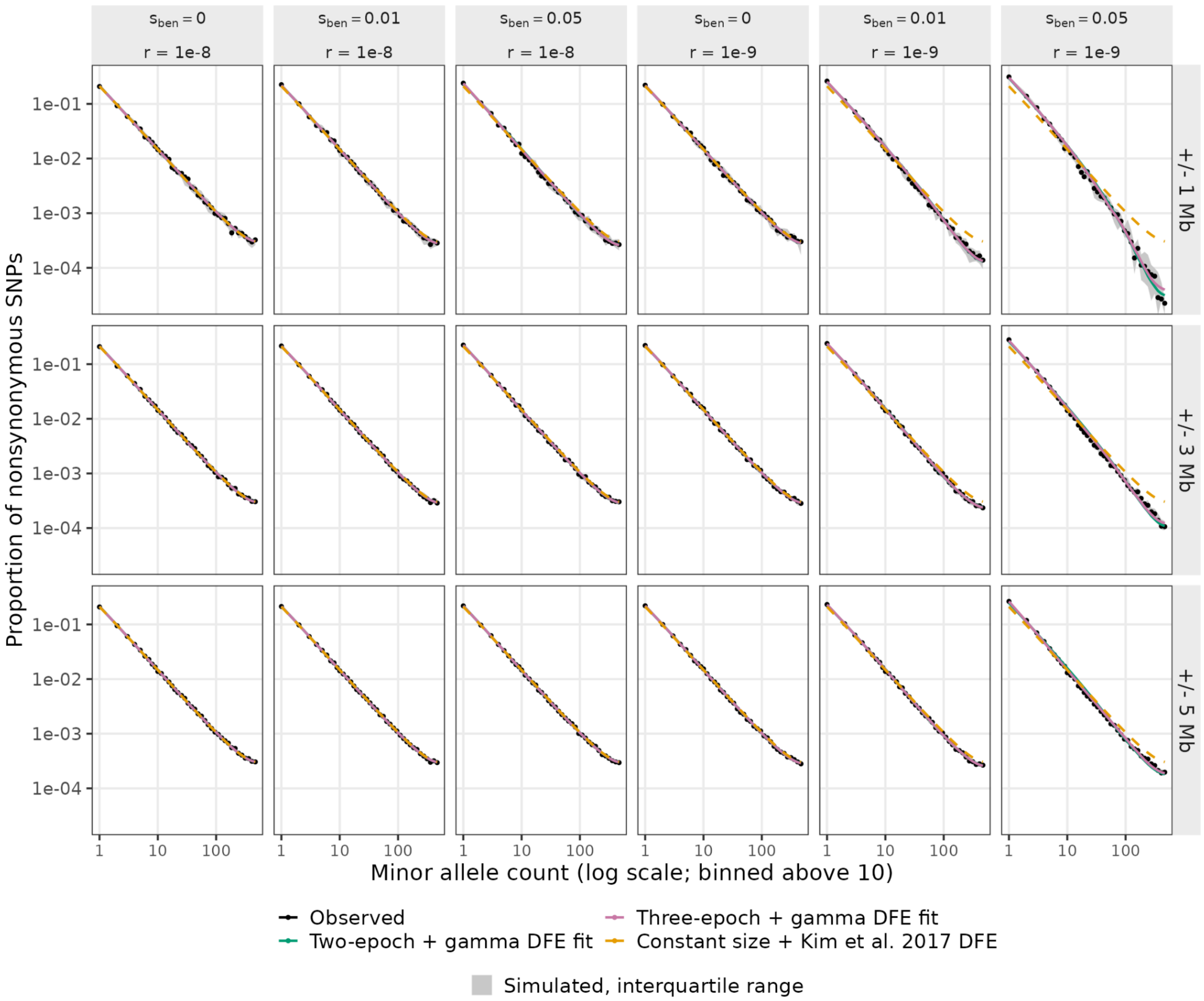

**Figure S10. Gamma DFE fits nonsynonymous SFS after a single completed sweep with *n* = 1000**

Folded nonsynonymous SFS ($n$ = 1000) from single-sweep simulations, shown as the proportion of nonsynonymous SNPs at each minor allele count (log scale). Columns show the sweep selection coefficient at each recombination rate. Rows show the window around the sweep used to build the SFS. Black points show the median observed proportion across 20 replicates, and grey shading shows the interquartile range across replicates. Green and pink lines show the median SFS from gamma DFEs fit conditional to a two-epoch and three-epoch demographic models, respectively. The dashed orange line shows the expected SFS under constant population size with the true human DFE (Kim et al. 2017).

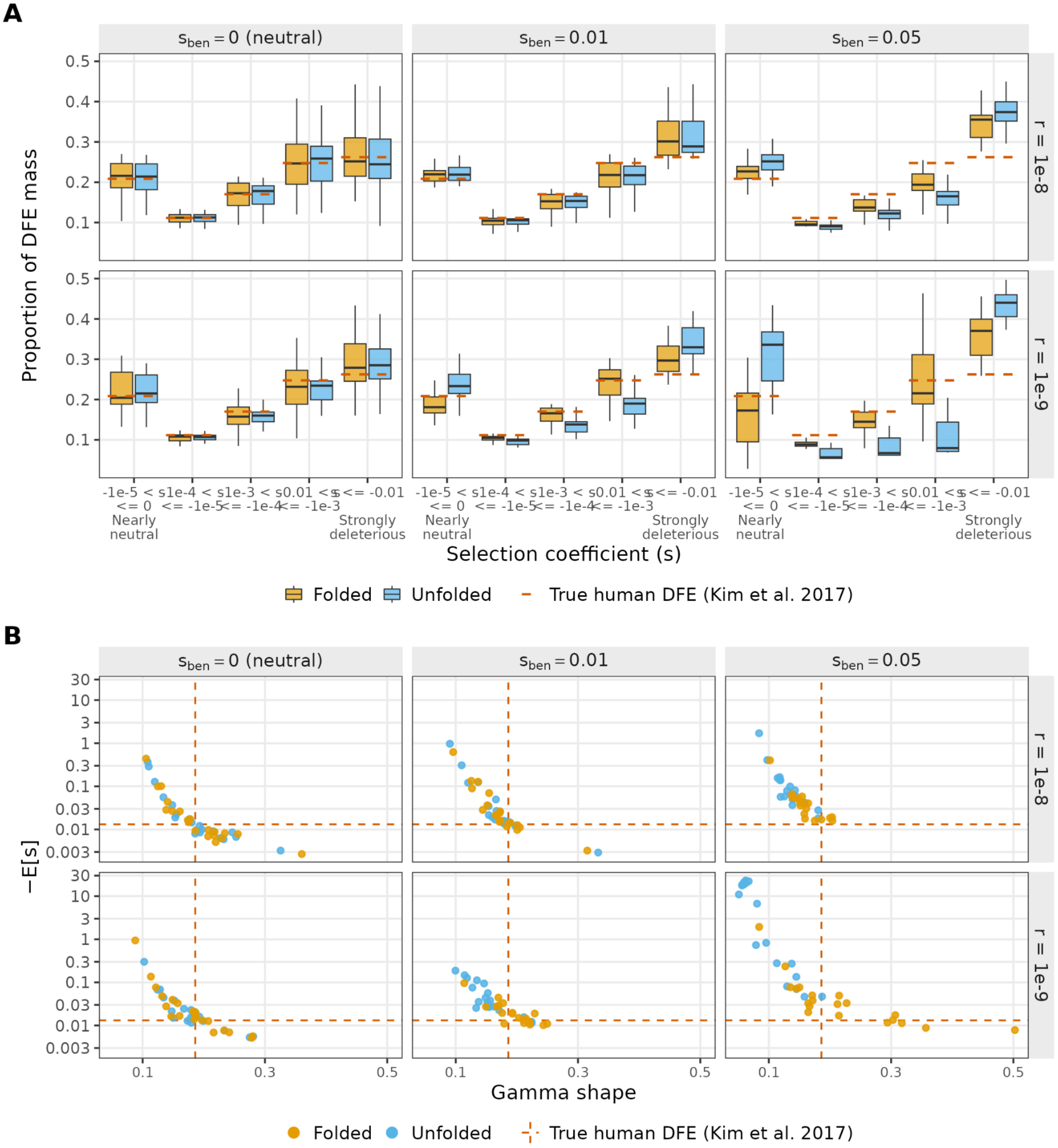

**Figure S11. Inference of the DFE using the unfolded SFS increases biases from a single completed selective sweep**

DFEs inferred after a single completed selective sweep using folded and unfolded SFS. Each gamma DFE was fitted from sites in windows $\pm\ 1Mb$ from the sweep center from sample sizes of $n$ = 100, conditional on the two-epoch demography fitted to the synonymous SFS. Columns show the beneficial mutation selection coefficient, $s_{ben}$, and rows show the recombination rate, *r*. **(A)** Proportion of DFE mass in each five selection coefficient bins. Boxes show median and interquartile range across 20 replicates, and whiskers extend 1.5x the interquartile range. **(B)** Inferred mean selection coefficient (-$E[s]$, log scale), versus the gamma shape parameter. Each point represents a replicate. In both panels, orange dashed lines mark the true human DFE (Kim et al. 2017; shape = 0.186, -$E[s]$ = 0.0131).

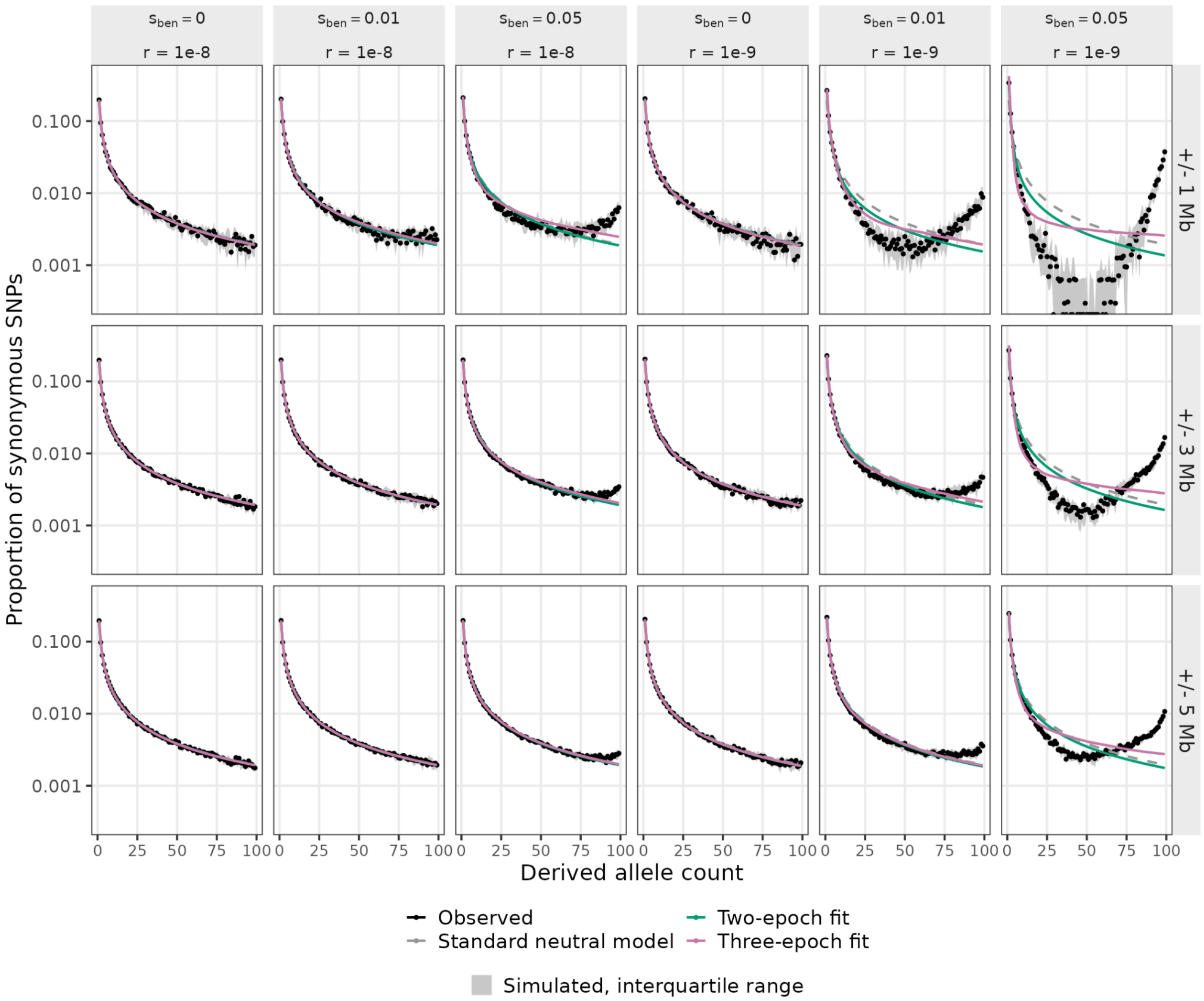

**Figure S12. Two- and three-epoch demographic models do not capture high frequency derived synonymous alleles in the unfolded SFS**

Unfolded synonymous SFS ($n$ = 100) from single-sweep simulations, shown as the proportion of synonymous SNPs at each derived allele count (log scale). Columns show the sweep selection coefficient at each recombination rate. Rows show the window around the sweep used to build the SFS. Black points show the median observed proportion across 20 replicates, and grey shading shows the interquartile range across replicates. The dashed grey line shows the standard neutral model. Green and pink lines show the median SFS from the two-epoch and three-epoch demographic models fitted to each replicate, respectively.

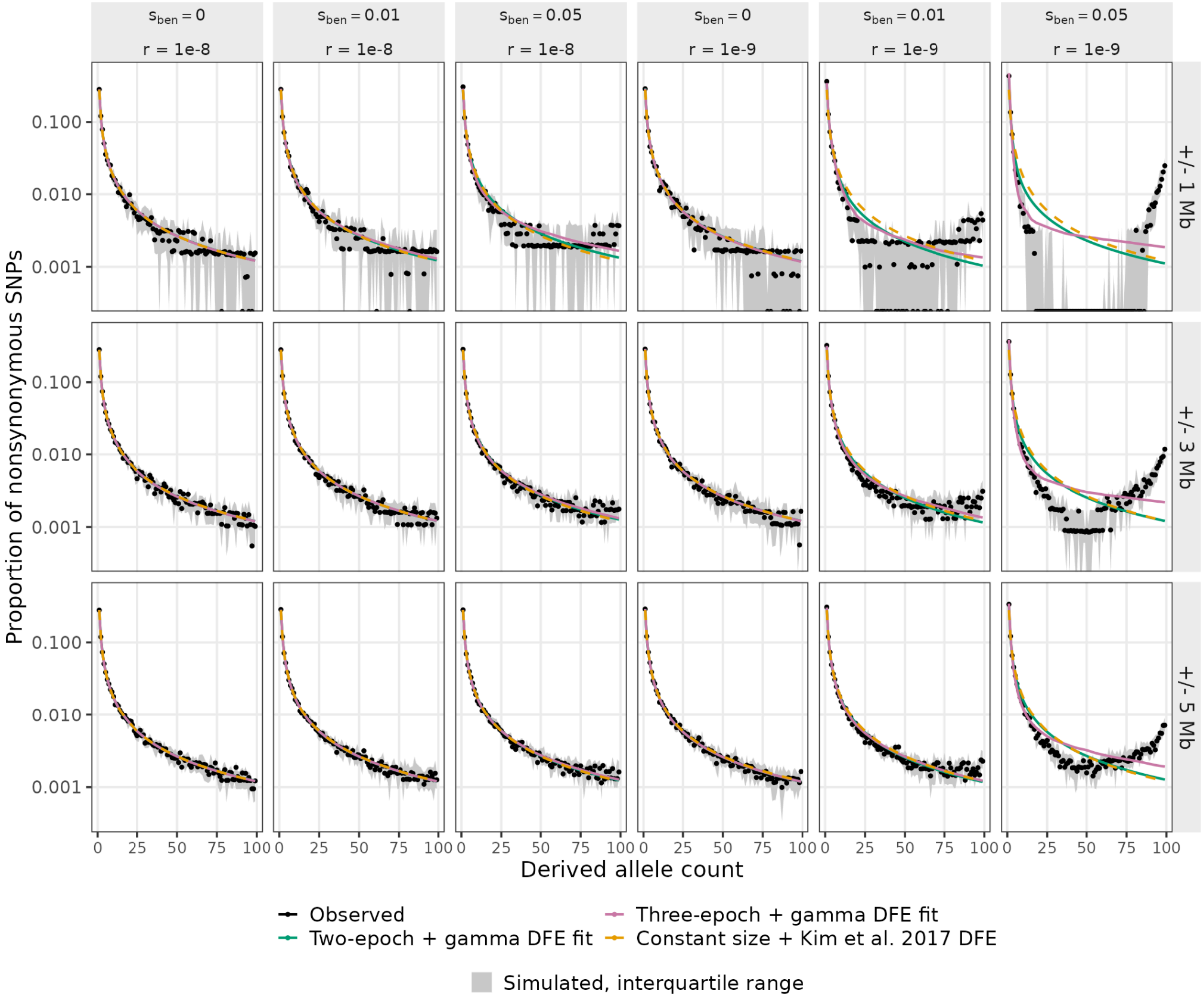

**Figure S13. The inferred DFE does not predict high frequency derived nonsynonymous alleles after a single completed sweep**

Unfolded nonsynonymous SFS ($n$ = 100) from single-sweep simulations, shown as the proportion of nonsynonymous SNPs at each derived allele count (log scale). Columns show the sweep selection coefficient at each recombination rate. Rows show the window around the sweep used to build the SFS. Black points show the median observed proportion across 20 replicates, and grey shading shows the interquartile range across replicates. Green and pink lines show the median SFS from gamma DFEs fit conditional to a two-epoch and three-epoch demographic models, respectively. The dashed orange line shows the expected SFS under constant population size with the true human DFE (Kim et al. 2017).

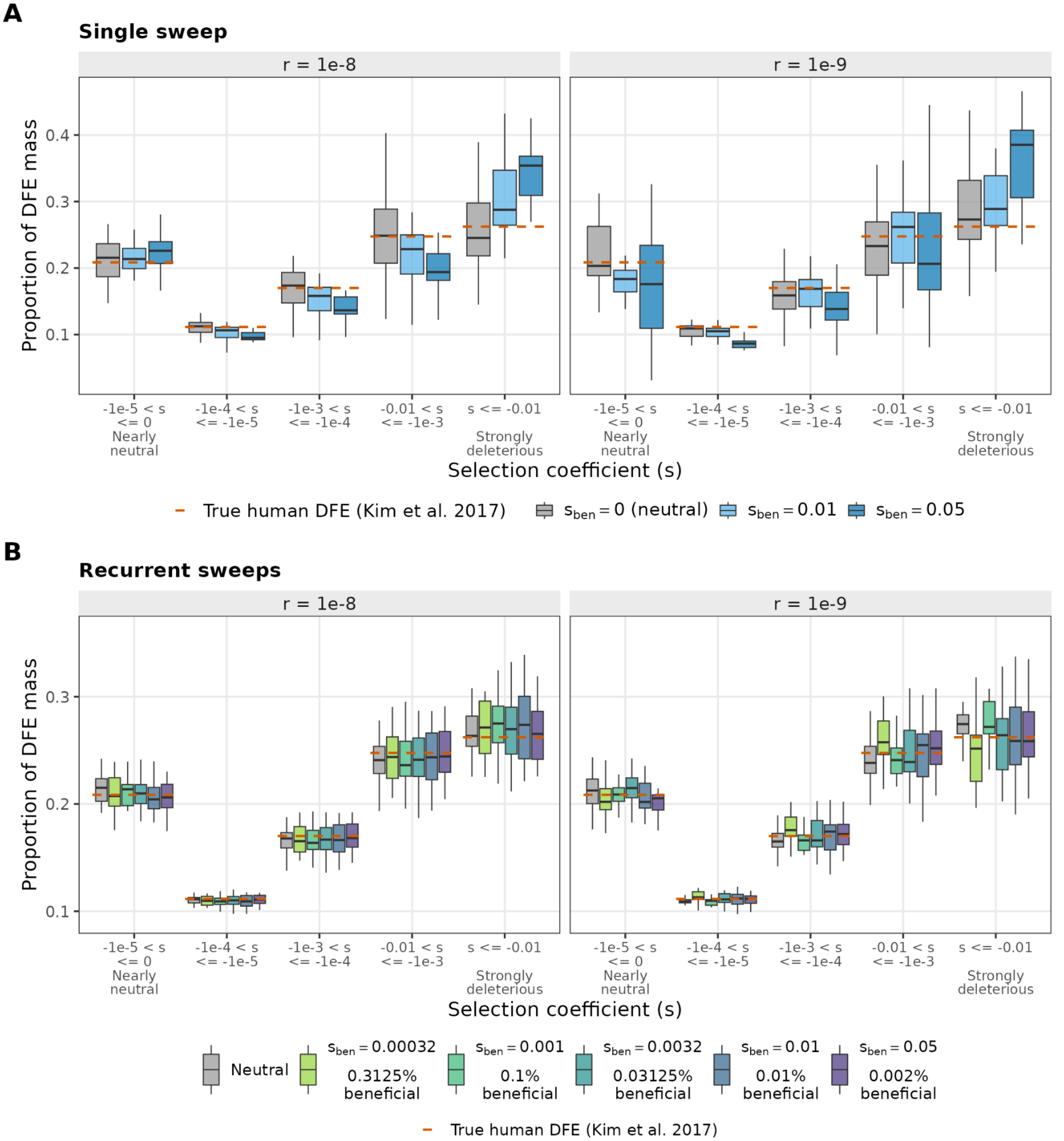

**Figure S14. Three-epoch demography gives similar DFE inferences as two-epoch demography**

Proportion of DFE mass in five selection coefficient bins, for gamma DFEs fitted to the folded nonsynonymous SFS ($n$ = 100), conditional on the three-epoch demographic model fitted to the synonymous SFS. Columns show recombination rate, $r$. **(A)** Single-sweep simulations from windows ±5 Mb from the center of the sweep. Colors represent the beneficial mutation selection coefficient, $s_{ben}$. **(B)** In each recurrent sweep condition, a fraction, *p*, of new nonsynonymous mutations is beneficial with selection coefficient, $s_{ben}$, where $log_{10}(s \times p) = -6$ and sites are drawn from the 10 Mb window. Boxes show the median and interquartile range across 20 replicates, and whiskers extend 1.5 x the interquartile range.

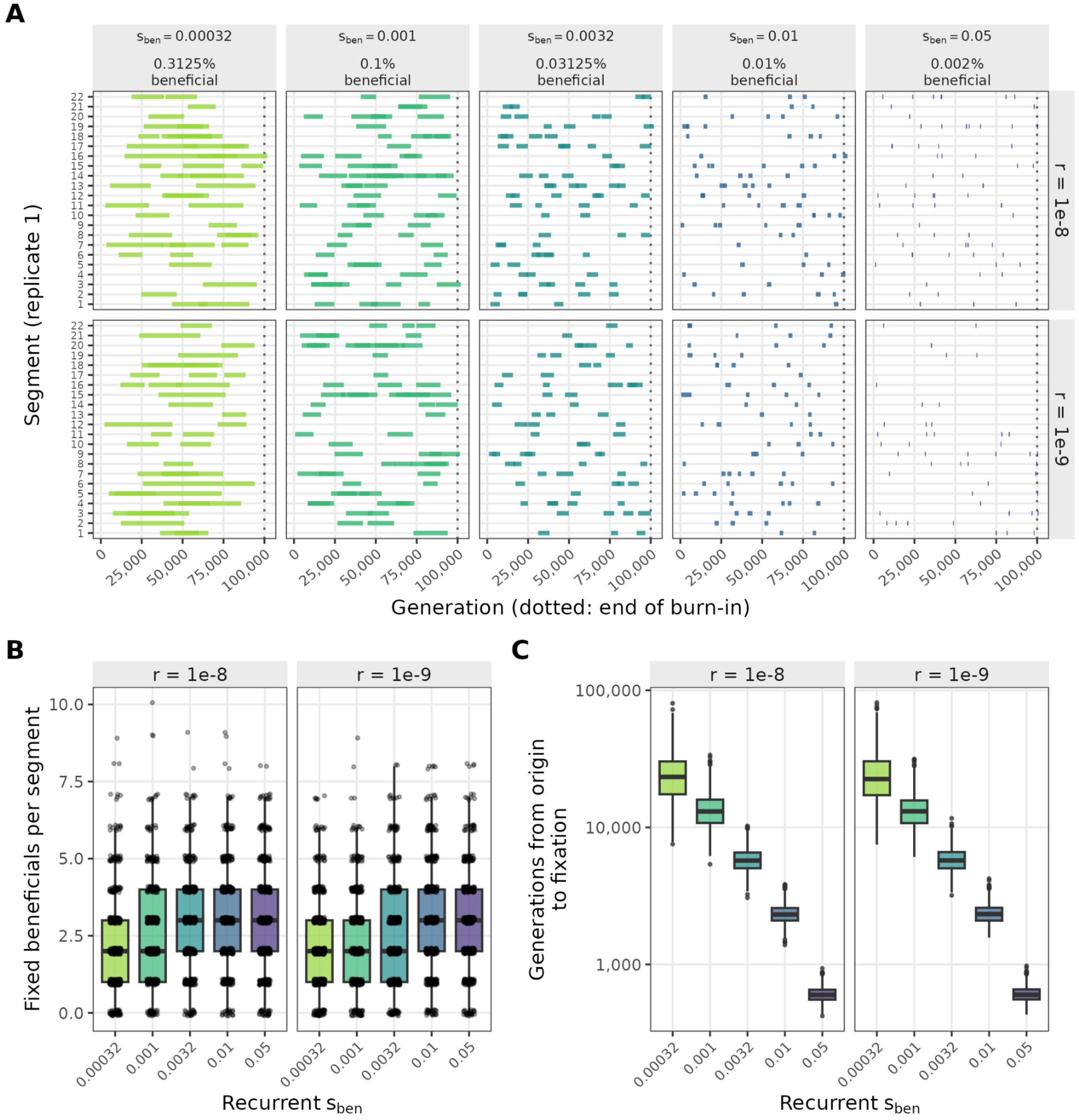

**Figure S15. Similar number of fixed beneficial mutations across selective strengths, but stronger sweeps fix in fewer generations**

Fixed beneficial mutations in the recurrent-sweep simulations. Beneficial mutations arise within a proportion of exonic nonsynonymous mutations (Table 1). Rows (A) and Columns (B, C) show recombination rate, *r*. **(A)** Generation of origin-to-fixation of each fixed beneficial mutation, shown as horizontal bar, for each simulated segment of replicate 1. **(B)** Number of fixed beneficial mutations fixed per segment. **(C)** Generations from origin to fixation (log scale) for fixed beneficial mutation. In B and C, boxes show median and interquartile range, whiskers extend 1.5x the interquartile range. In B, points are shown for each individual segment. In C, points show values that extend past whiskers.

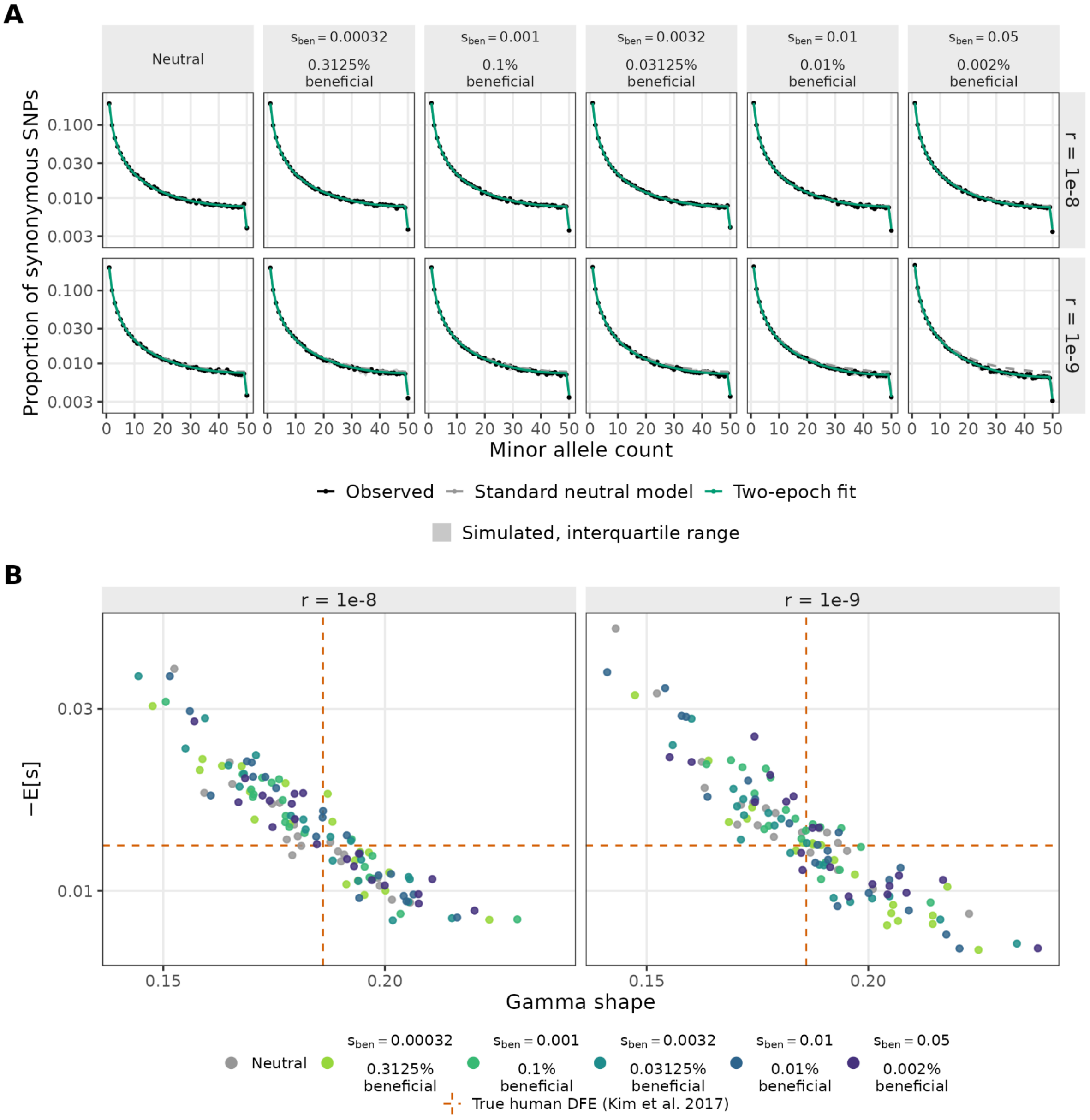

#### Figure S16. Recurrent selective sweeps do not influence demographic and DFE inferences

Recurrent-sweep inferences using the folded SFS ($n$ = 100). In each condition, a fraction, $p$, of new nonsynonymous mutations is beneficial with selection coefficient, $s_{ben}$, where $log_{10}(s \times p) = -6$ and sites are drawn from the 10 Mb window. Neutral controls use the same window. **(A)** Synonymous SFS shown as the proportion of synonymous SNPs at each minor allele count (log scale). Columns show recurrent sweep condition and rows show recombination rate. Black points show the median observed proportion across replicates, and shading shows the interquartile range of simulated spectra. The green line is the two-epoch fit and the grey dashed line is the expected SFS under the standard neutral model. Both are the median across replicates. **(B)** Inferred -E[s] (log-scale) plotted against gamma shape for gamma DFEs fit to the nonsynonymous SFS, conditional on a two-epoch model. Each point is one replicate, color represents the condition. Recombination is denoted by row in A and by column in B. In both panels, dashed orange lines represent the true simulated human DFE (Kim et al. 2017); gamma shape = 0.186, E[s] = -0.0131.

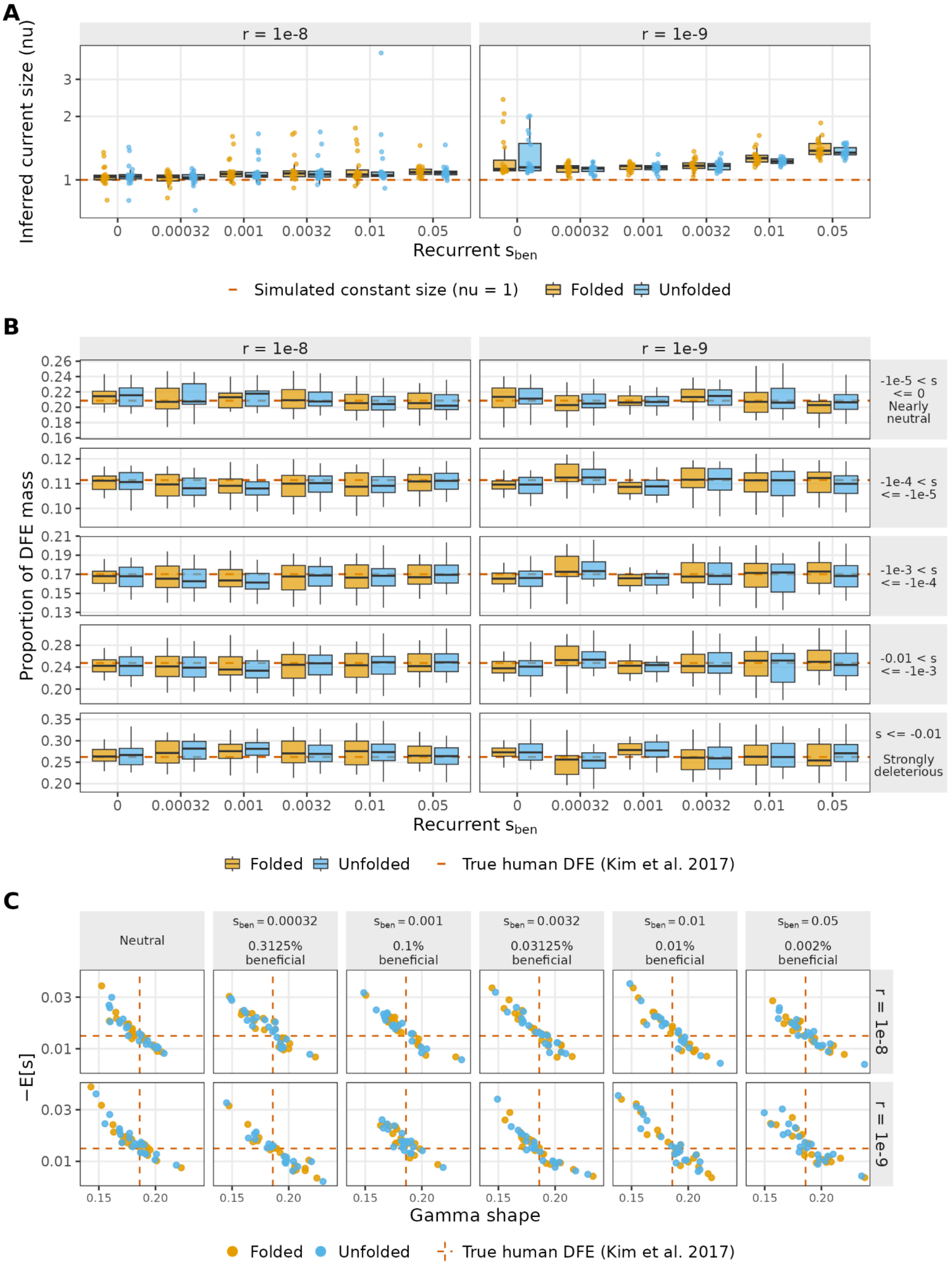

A
r = 1e-8
r = 1e-9
Inferred current size (nu)
Recurrent $s_{ben}$
Simulated constant size (nu = 1)
Folded
Unfolded
B
Proportion of DFE mass
-1e-5 < s <= 0 Nearly neutral
-1e-4 < s <= -1e-5
-1e-3 < s <= -1e-4
-0.01 < s <= -1e-3
s <= -0.01 Strongly deleterious
True human DFE (Kim et al. 2017)
C
Neutral
$s_{ben} = 0.00032$ 0.3125% beneficial
$s_{ben} = 0.001$ 0.1% beneficial
$s_{ben} = 0.0032$ 0.03125% beneficial
$s_{ben} = 0.01$ 0.01% beneficial
$s_{ben} = 0.05$ 0.002% beneficial
−E[s]
Gamma shape

#### Figure S17. In recurrent sweep simulations, folded and unfolded SFS yield similar demography and DFE inferences

Two-epoch demography and gamma DFE inferences from recurrent-sweep simulations using the folded or unfolded SFS (*n*=100) and sites drawn from the full 10 Mb segment. Unfolded SFS summarizes the derived allele count whereas folded summarizes the minor allele count. In each condition, a fraction, *p*, of new nonsynonymous mutations is beneficial with selection coefficient, $s_{ben}$, where $log_{10}(s \times p) = -6$. Color represents folding. Dashed orange lines represent the true simulated human DFE (Kim et al. 2017); gamma shape = 0.186, E[s] = -0.0131. **(A)** Current population size (ν, log scale) inferred by the two-epoch demographic model fitted to the synonymous SFS. Recurrent sweep condition is along the x-axis. **(B)** Proportion of DFE mass in five selection coefficient bins, conditional on the two-epoch demographic model fitted to the synonymous SFS. Rows represent recurrent sweep conditions. **(C)** Inferred -E[s] (log-scale) plotted against gamma shape for gamma DFEs fit to the nonsynonymous SFS, conditional on a two-epoch model. Each point is one replicate. Columns represent recurrent sweep conditions. Recombination rate, *r*, is denoted in columns in A and B and rows in C.

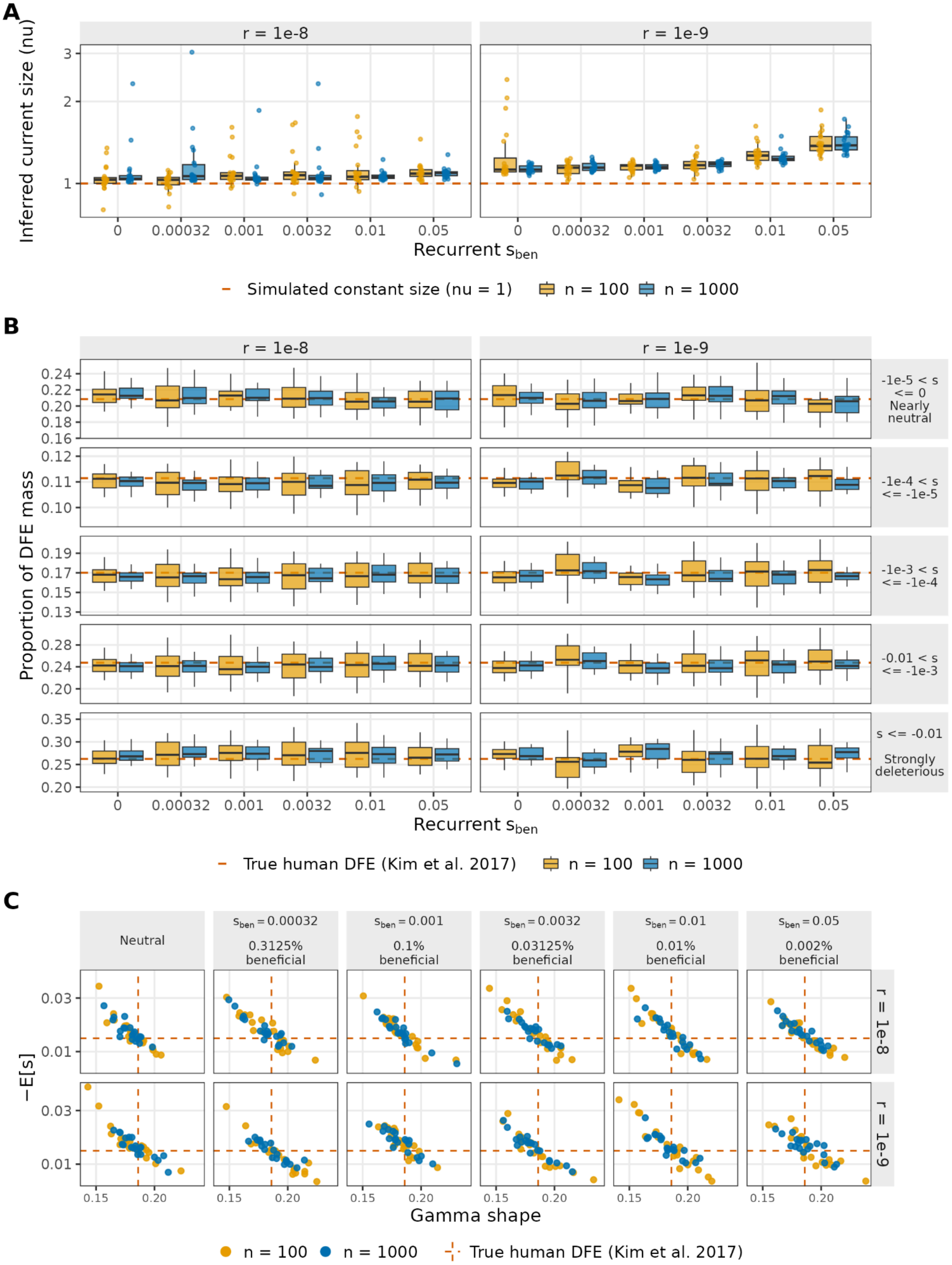

A
r = 1e-8
r = 1e-9
Inferred current size (nu)
3
2
1
0
0.00032
0.001
0.0032
0.01
0.05
Recurrent $s_{ben}$
Simulated constant size (nu = 1)
n = 100
n = 1000
B
Proportion of DFE mass
-1e-5 < s <= 0 Nearly neutral
-1e-4 < s <= -1e-5
-1e-3 < s <= -1e-4
-0.01 < s <= -1e-3
s <= -0.01 Strongly deleterious
True human DFE (Kim et al. 2017)
C
Neutral
$s_{ben}$ = 0.00032 0.3125% beneficial
$s_{ben}$ = 0.001 0.1% beneficial
$s_{ben}$ = 0.0032 0.03125% beneficial
$s_{ben}$ = 0.01 0.01% beneficial
$s_{ben}$ = 0.05 0.002% beneficial
−E[s]
Gamma shape

**Figure S18. DFE inferences are similar between sample sizes in recurrent sweep simulations**

Two-epoch demography and gamma DFE inferences from recurrent-sweep simulations using sample sizes of $n$ = 100 and $n$ = 1000, folded SFS, and sites drawn from the full 10 Mb segment. In each condition, a fraction, $p$, of new nonsynonymous mutations is beneficial with selection coefficient, $s_{ben}$, where $log_{10}(s \times p) = -6$. Color represents sample size. Dashed orange lines represent the true simulated human DFE (Kim et al. 2017); gamma shape = 0.186, E[s] = -0.0131. **(A)** Current population size (ν, log scale) inferred by the two-epoch demographic model fitted to the synonymous SFS. Recurrent sweep condition is along the x-axis. **(B)** Proportion of DFE mass in five selection coefficient bins, conditional on the two-epoch demographic model fitted to the synonymous SFS. Rows represent recurrent sweep conditions. **(C)** Inferred -E[s] (log-scale) plotted against gamma shape for gamma DFEs fit to the nonsynonymous SFS, conditional on a two-epoch model. Each point is one replicate. Columns represent recurrent sweep conditions. Recombination rate, $r$, is denoted in columns in A and B and rows in C.

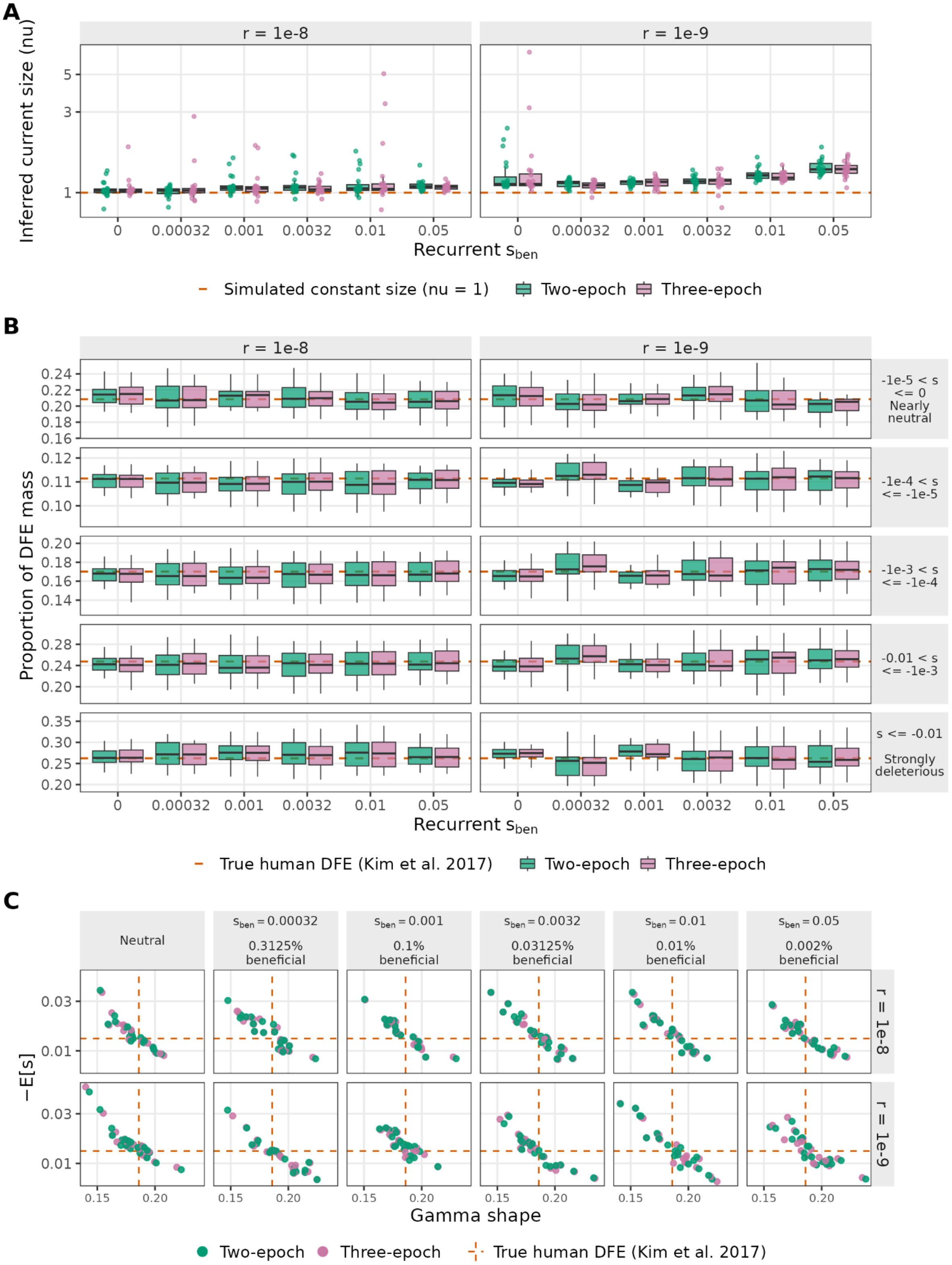
A
r = 1e-8
r = 1e-9
Inferred current size (nu)
Recurrent $s_{ben}$
Simulated constant size (nu = 1)
Two-epoch
Three-epoch
B
Proportion of DFE mass
-1e-5 < s <= 0 Nearly neutral
-1e-4 < s <= -1e-5
-1e-3 < s <= -1e-4
-0.01 < s <= -1e-3
s <= -0.01 Strongly deleterious
True human DFE (Kim et al. 2017)
C
Neutral
$s_{ben} = 0.00032$ 0.3125% beneficial
$s_{ben} = 0.001$ 0.1% beneficial
$s_{ben} = 0.0032$ 0.03125% beneficial
$s_{ben} = 0.01$ 0.01% beneficial
$s_{ben} = 0.05$ 0.002% beneficial
−E[s]
Gamma shape

**Figure S19. DFE inferences are similar after conditioning on two- or three-epoch demographic models in recurrent sweep simulations**

Two-epoch demography and gamma DFE inferences from recurrent-sweep simulations using two- and three-epoch demographic models, folded SFS, and sites drawn from the full 10 Mb segment. In each condition, a fraction, *p*, of new nonsynonymous mutations is beneficial with selection coefficient, $s_{ben}$, where $log_{10}(s \times p) = -6$. Color represents demographic models. Dashed orange lines represent the true simulated human DFE (Kim et al. 2017); gamma shape = 0.186, E[s] = -0.0131. **(A)** Current population size (ν, log scale) inferred by the two-epoch demographic model fitted to the synonymous SFS. Recurrent sweep condition is along the x-axis. **(B)** Proportion of DFE mass in five selection coefficient bins, conditional on the two-epoch demographic model fitted to the synonymous SFS. Rows represent recurrent sweep conditions. **(C)** Inferred -E[s] (log-scale) plotted against gamma shape for gamma DFEs fit to the nonsynonymous SFS, conditional on the respective model. Each point is one replicate. Columns represent recurrent sweep conditions. Recombination rate, *r*, is denoted in columns in A and B and rows in C.

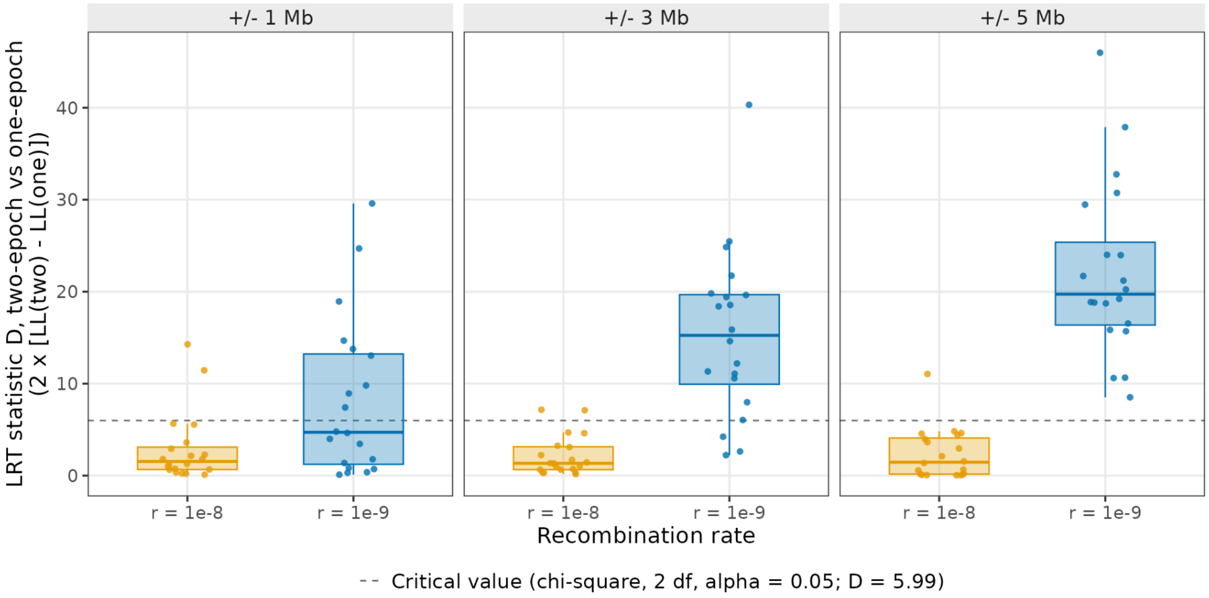

**Figure S20. Without a sweep, a two-epoch demographic model fits the synonymous SFS better at low recombination rates**

Likelihood-ratio test comparing the two-epoch and one-epoch demographic model, fitted to the folded synonymous SFS ($n$ = 100) in simulations without a sweep. Panels show the window used to build the SFS and the x-axis shows the recombination rate, *r*. The test statistic is $D = 2 \times [LL_2 - LL_1]$. The dashed line marks the $\chi^2$ critical value (2 df, $\alpha = 0.05$, D = 5.99). Boxes show the median and interquartile range across 20 replicates, whiskers extend 1.5 x IQR, and points show individual replicates. All replicates converged.
